# Molecular determinants underlying substrate receptor specificity of human CRL4B E3 ubiquitin ligase

**DOI:** 10.64898/2026.03.13.711546

**Authors:** Weaam I. Mohamed, Julius Rabl, Bilal M. Qureshi, Alexander Leitner, Federico Uliana, Arghavan Kazemzadeh, Norhan Radwan, Matthias Peter

**Affiliations:** Institute of Biochemistry, Department of Biology, ETH Zürich, Otto-Stern-Weg 3, 8093 Zürich, Switzerland; Cryo-EM Knowledge Hub (CEMK), Otto-Stern-Weg 3, 8093 Zürich, Switzerland; Scientific Center for Optical and Electron Microscopy (ScopeM), ETH Zurich, Otto-Stern-Weg 3, 8093 Zürich, Switzerland; Institute of Molecular Systems Biology, Department of Biology, ETH Zürich, Otto-Stern-Weg 3, 8093 Zürich, Switzerland; Biozentrum II, Johannes Gutenberg University Mainz, Hanns-Dieter Hüsch-Weg 17, 55128 Mainz, Germany; Department of Chemistry and Applied Biosciences, ETH Zürich, Vladimir-Prelog-Weg 1-5/10, 8093 Zürich, Switzerland

## Abstract

The vertebrate CRL4 family of Cullin-RING E3 ubiquitin ligases is distinguished from other cullin-based ligases by the presence of two highly homologous paralogs, CUL4A and CUL4B. Both CRL4 complexes use the DDB1 subunit to recruit dedicated and interchangeable substrate receptors called DCAFs, but the underlying mechanisms guiding DCAF specificity for CUL4B or CUL4A remain poorly understood. Here, we performed structural and biochemical analyses of the CRL4B^LIS1^ complex and identified two molecular determinants for CUL4B-specific DCAFs. First, we discovered that the unique CUL4B N-terminal extension directly binds CUL4B-specific DCAFs, enhancing their complex formation. This direct interaction can be modulated by phosphorylation, adding the possibility for spatiotemporal regulation. Second, the cryo-EM model of the CRL4B^LIS1^ complex identified a novel interface on the DDB1 subunit which promotes LIS1 recruitment. Quantitative affinity measurements and mutational analysis confirmed that this DDB1 interface is generally important for recruiting CUL4B-specific DCAFs including WDR1 and BRWD1, but not for CUL4A-specifc DCAFs, such as DCAF8. Together, our study identifies molecular determinants and unexpected interfaces on CRL4 components that dictate preference for DCAF recruitment.

**Graphical Abstract showing how the two CRL4 complexe s, namely CRL4A and CRL4B, recruit their DCAFs:** 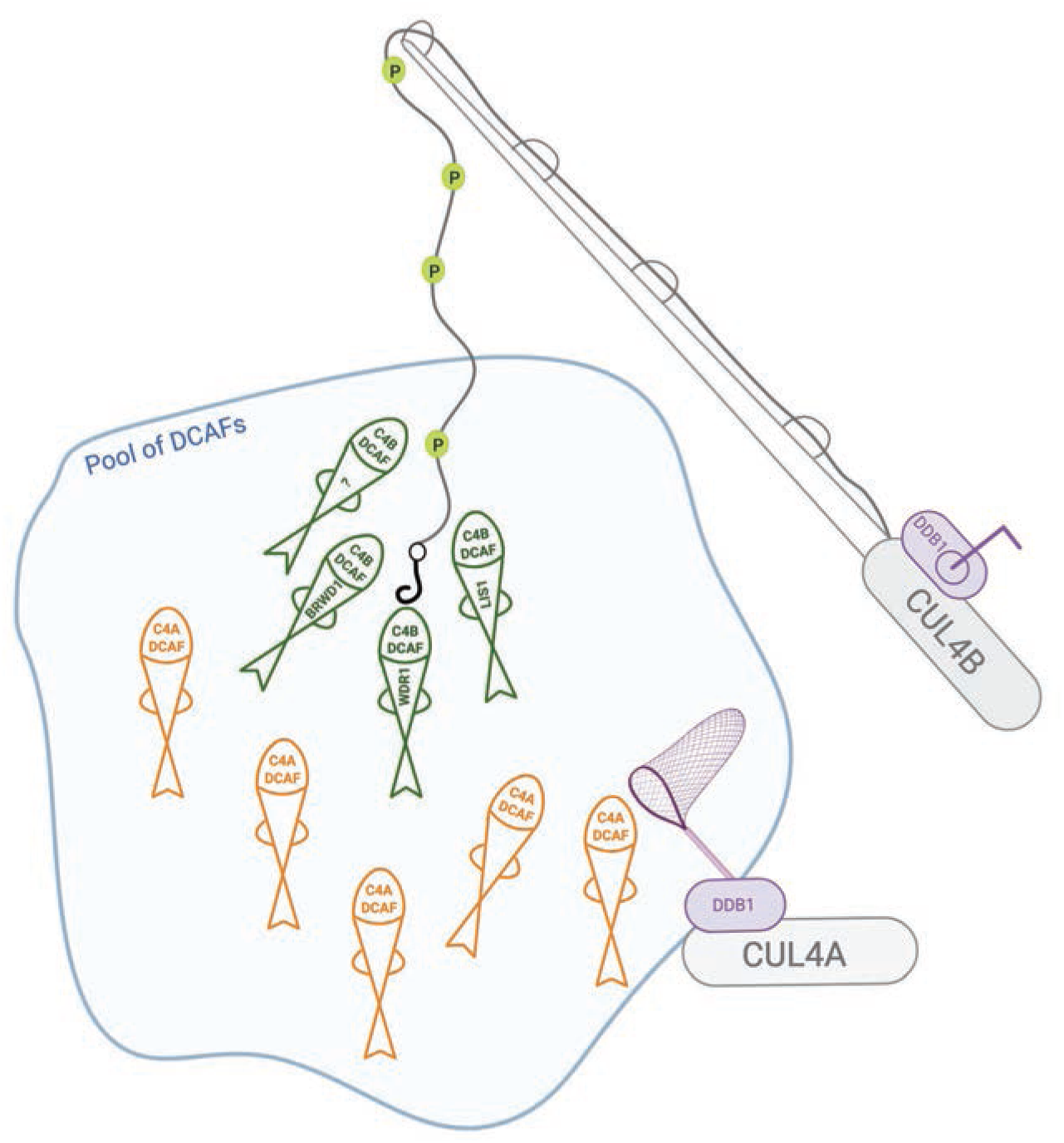

## Introduction

The last step of the ubiquitination process, covalently attaching ubiquitin to substrates, is performed by E3 ligases. They confer specificity of the ubiquitination machinery and regulate localized and temporal substrate ubiquitination (Deshaies & Joazeiro, 2009), altering the activity and/or stability of target proteins. Cullin RING ubiquitin ligases (CRL) comprise the largest family of E3 ligases, owing to their modular assembly with multiple substrate receptors, encompassing approximately 250 members.

The CRL4 family is unique among CRLs, with an evolutionary divergence in which yeast, worms, plants and flies possess only one CUL4 gene, while vertebrates evolved two structurally similar paralogs, CUL4A and CUL4B. The paralogs are distinguished by a unique CUL4B amino terminal extension, and they exhibit overlapping and distinct cytoplasmic and nuclear functions (Nakagawa & Nakagawa, 2025). Disruption of the *CUL4* gene in yeast leads to mitotic defects and hypersensitivity to DNA damage, while loss of CUL4 activity in *C. elegans*, fruit flies, and zebrafish, leads to severe developmental defects (Holmberg et al., 2005; Nakagawa & Nakagawa, 2025; Zhao et al., 2010). In contrast, CUL4A-knockout mice do not show apparent developmental defects except male infertility and cardiac hypertrophy (Kopanja et al., 2011). However, deletion of CUL4B in mice is embryonically lethal, characterized by significant apoptosis in extra-embryonic tissues (Liu et al., 2009). If CUL4B is only disrupted in mouse epiblasts, CUL4B-deficient embryos can survive but they exhibit brain development defects (Jiang et al., 2012; Liu et al., 2012). This phenotype resembles CUL4B loss-of-function mutants in human patients suffering from X-linked intellectual disability (XLID) (Tarpey et al., 2007, 2009; Zou et al., 2007). To-date, the molecular and structural distinctions of CUL4A and CUL4B E3 ligase complexes are poorly understood.

The CUL4 scaffold recruits DNA Damage-Binding protein 1 (DDB1) at its N-terminal Cullin Repeat Domain 1 (CRD1) and RING Box protein 1 (RBX1) at the C-terminus (CTD) to assemble CRL4 (CUL4-DDB1-RBX1) complexes (Angers et al., 2006). RBX1 is a RING domain containing protein directly involved in catalysis by binding ubiquitin-loaded E2 ubiquitin-conjugating enzymes. In human cells, approximately 90% of CUL4 is in complex with DDB1 (Reichermeier et al., 2020), which in turn recruits dedicated substrate receptors called DDB1 Cullin4 Associated Factors (DCAFs). DDB1 is composed of three β-propeller domains A, B and C (BPA, BPB, BPC). While BPB interacts with the CUL4 scaffold, BPA and BPC together form a cleft, which is essential for binding to the helix-loop-helix (HLH) or HLH-like motif within DCAFs (Figure 1A) (Angers et al., 2006). In addition, DDB1 binding of some DCAFs such as DCAF15, Cockayne syndrome WD repeat protein (CSA), and DCAF5 is dependent on DET1 DDB1 Associated protein 1 (DDA1) (Burgess et al., 2025; Bussiere et al., 2020; Llerena Schiffmacher et al., 2024; Radko-Juettner et al., 2024), first identified as a subunit of a plant CRL4 complex (Lau et al., 2011; Zhang et al., 2008). Interestingly, DDA1 docks to the bottom surface of the DDB1 BPA domain, which is far removed from both the CUL4A- and DCAF-binding sites. The majority of DCAFs contain WD40 domains, which supports DDB1 binding, but also enhances substrate recruitment (Lee & Zhou, 2007). Conventional DCAFs do not directly interact with the CUL4 scaffold and most DCAFs cannot distinguish CUL4A and CUL4B. However, a subset exhibits preferential binding to CUL4A or CUL4B (Reichermeier et al., 2020). For example, Cereblon (CRBN) and DCAF8 prefer CUL4A, while PH-interacting protein (PHIP), Bromodomain and WD repeat-containing protein 1 (BRWD1) and DCAF10 exhibit higher affinity towards CUL4B (Reichermeier et al., 2020). Importantly, CUL4B also binds non-canonical DCAFs, such as the mitotic-specific substrate receptors Lissencephaly 1 (LIS1) and WD repeat-containing protein 1 (WDR1) (Stier et al., 2023). Their cell cycle regulation is mediated by phosphorylation of the unique CUL4B N-terminus during mitosis, which increases binding of LIS1 and WDR1. This regulation is lost in the XLID patient-derived P50L mutant, due to decreased phosphorylation of T49 in mitosis (Stier et al., 2023). Likewise, patients suffering from lissencephaly exhibit heterozygous mutations in LIS1, and protein levels of mutated LIS1 correlate with clinical severity (Zillich et al., 2025).

**Figure 1:**
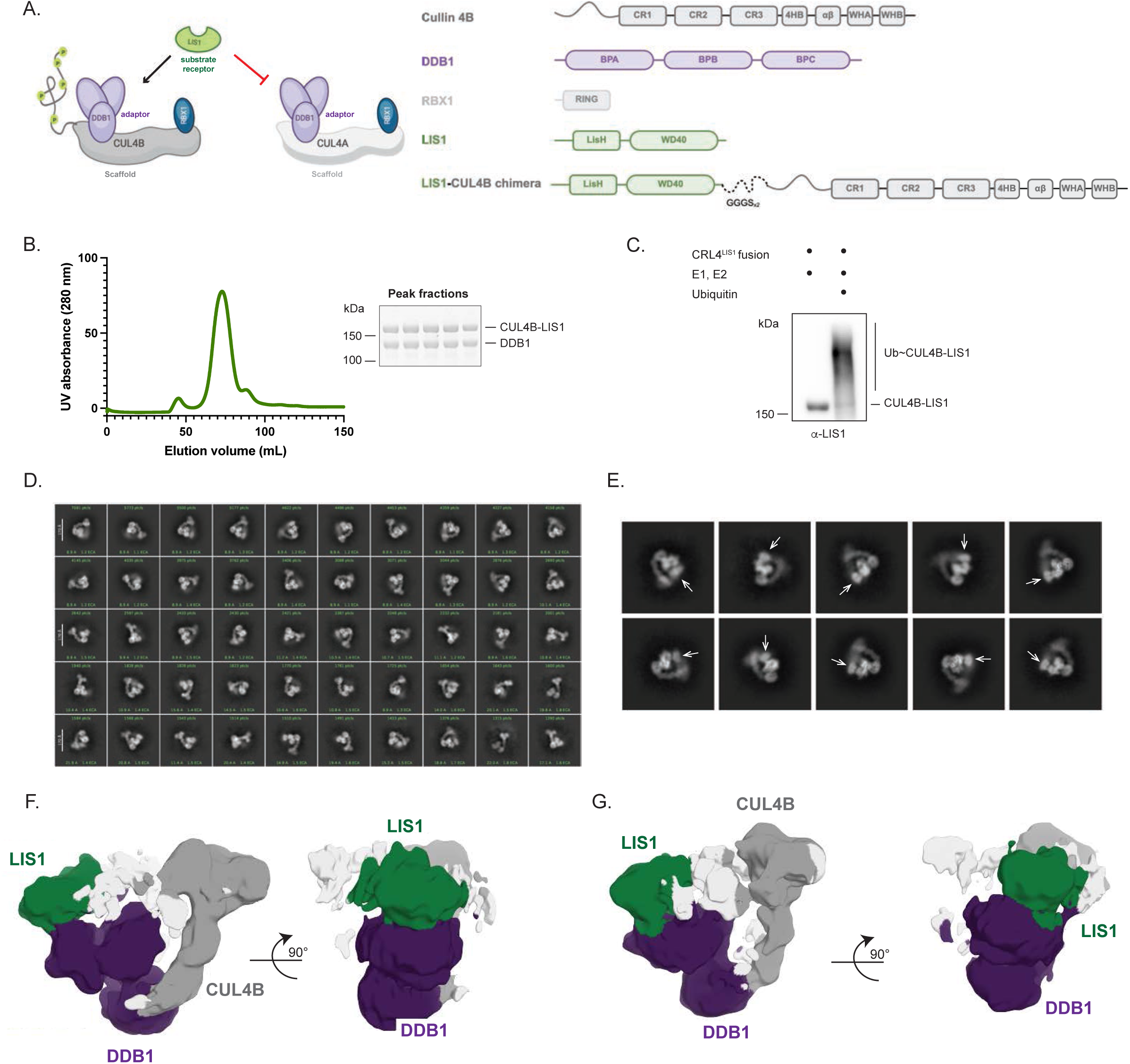
Biochemical and structural analysis of the CRL4B^LIS1^ complex. **A.** Schematic drawing of CUL4A and CUL4B complexes (left), which discriminate binding of the substrate-receptor LIS1. The phosphorylated N-terminal extension of CUL4B is indicated. The structural domain arrangement within each of the CRL4B^LIS1^ subunits is shown (right). The LIS1-CUL4B chimera protein is produced with a 10-amino acid linker separating the two proteins. CR1, 2, and 3: Cullin Repeat 1,2, and 3; 4HB: 4 Helix-Bundle; α/β: alpha/beta domain; WH1, WH2: Wing-helix domains 1 and 2. **B.** The elution profile of absorbance (280 nm) per fraction (ml) of recombinant CRL4B^LIS1^ complexes was analyzed by size exclusion chromatography, revealing a major peak of the CUL4B-LIS1 chimera bound to DDB1. The stoichiometric complex present in the peak fractions was analyzed by SDS-PAGE and Coomassie-staining (right). **C.** Auto-ubiquitination assay of CRL4B^LIS1^ complexes incubated together with the ubiquitination machinery (E1, E2, MgCl_2_, and ATP), in the presence or absence of ubiquitin (Ub). The reactions were analyzed by immunoblotting with LIS1 antibodies. The position of the unmodified CUL4B-LIS1 chimera and ubiquitinated species (Ub-CUL4B-LIS1) are indicated. **D.** and **E**. 2D classification of CRL4B^LIS1^ particles using CryoSparc software (left). Scale bars: 170 Å. A zoom-in view of representative CRL4B^LIS1^ complexes showing a potential LIS1 density (**E**). The white arrow points at the extra density on top of DDB1, which corresponds to the LIS1 propeller. **F.** and **G.** 3D reconstruction of the two major classes that contain fully assembled CRL4B^LIS1^ complexes. The maps have been aligned with respect to DDB1 (purple). The position of CUL4B (grey) and LIS1 (green) differs between the extended (**F**) and the closed conformation (**G**). In the closed conformation, LIS1 and the C-terminus of CUL4B are in direct proximity. Additional density (white) is observed but cannot be assigned due to limited resolution of the maps.

LIS1 is a regulatory co-factor for the assembly of Dynein-Dynactin complex, regulating microtubule dynamics and spindle positioning in mitosis. Indeed, cells lacking LIS1 or CUL4B function exhibit misplaced spindles, suggesting that a mitotic CUL4B^LIS1^ E3 ligase complex ubiquitinates substrates regulating dynein-dynactin activity (Singh et al., 2024). LIS1 contains a WD40 domain and a LISH motif, which promotes homodimerization (Mohamed et al., 2021) (PDB 1UUJ, PDB 6IWV). Moreover, LIS1 encompasses a putative HLH motif that may contribute to DDB1 binding (Stier et al., 2023). However, the molecular mechanisms explaining how LIS1 and other DCAFs specifically interact with CUL4B remain obscure.

In this study we used a structural approach visualizing CRL4B^LIS1^ complexes to unravel common determinants distinguishing CUL4B and CUL4A substrate receptors. Surprisingly, our cryo-EM model shows that the WD40 domain of LIS1 contacts the BPA domain of DDB1, while canonical DCAFs bind to the BPC domain. In addition, LIS1 directly binds the CUL4B N-terminal extension, and this interaction is promoted by phosphorylation. These novel specificity determinants are also used by WDR1 and BRWD1, implying that they generally distinguish CUL4A and CUL4B specific DCAFs.

## Results

### Structural analysis of fully assembled CRL4B^LIS1^ complexes

We previously reported that the CRL4B-specific DCAFs LIS1 and WDR1 bind to CUL4B with almost 10-fold higher affinity compared to CUL4A (Figure 1A) (Stier et al., 2023). In order to understand the underlying specificity mechanism, we performed a structural analysis of the reconstituted CRL4B^LIS1^ complex using Cryo-EM. To promote stochiometric and stable complex formation, we fused the LIS1 protein to CUL4B via a flexible linker of 10-amino acids (GGGGSx2) (Figure 1A). The chimeric CUL4B-LIS1 fusion protein was soluble and readily assembled a stoichiometric complex with RBX1 and DDB1 as shown by gel filtration (Figure 1B). Moreover, the chimeric CRL4B^LIS1^ was catalytically active and was readily auto-ubiquitinated *in vitro* (Figure 1C), implying that the CUL4B-LIS1 fusion protein folds into its native functional state.

We studied the chimeric CRL4B^LIS1^ complex (CUL4B-LIS1, RBX1, DDB1) by single-particle cryo-EM analysis (Supp. Figure 1). The detected particles were classified into three major cohorts: 1) DDB1 alone, 2) CUL4B-DDB1, and 3) CUL4B-DDB1 including an extra EM density above DDB1 (Figure 1D and E) consistent with LIS1 binding. These DDB1-CUL4B-LIS1 particles displayed conformational heterogeneity, as the CUL4B arm can adopt different positions relative to DDB1 ranging from an extended conformation (Figure 1F) to a closed conformation which brings the C-terminus of CUL4B and LIS1 into direct proximity (Figure 1G). A focused refinement of DDB1-LIS1 resulted in a map with an overall resolution (FSC 0.143) of 4.3 Å, with the resolution of the LIS1 region of the map was 7-10 Å (Supp. Figure 1).

While structural heterogeneity precluded atomic resolution, the map was of sufficient quality to allow definite placement of LIS1 and DDB1 into the density.

## Identification of a conserved DDB1 REKE loop at the interface with LIS1

Interestingly, the CRL4B^LIS1^ cryo-EM map indicated a novel interaction surface between the WD40 domain of LIS1 and the BPA of DDB1 (Figure 2A). Using a mask around these DDB1 and LIS1 domains, followed by several focused refinement rounds, the resolution of this interaction interface could be resolved to ca. 7 Å (Figure 2A, Supp. Figure 1), which allowed docking of the LIS1 model into the map by rigid-body fitting. The interaction between LIS1 and DDB1 appears mostly mediated by salt bridges between R111, K153, R158, R198, and E201 of DDB1 and R212, E235, E276, and D338 of LIS1. Moreover, W340 likely contributes to the interaction by π-stacking with arginine residues of DDB1 (Figure 2C). Interestingly, this analysis highlighted a loop within the DDB1 BPA, in accordance with its amino acid sequence hereafter called REKE loop, near the LIS1 WD40 domain (Figure 2B). Due to the limited resolution of the CRL4B^LIS1^ cryo-EM map, we were unable to resolve the corresponding LIS1 residues interacting with the REKE loop. Likewise, we could not visualize the conserved HLH motif located in the N-terminus of LIS1 (Figure 2C), which is required for its interaction with DDB1 (Stier et al., 2023). While we cannot exclude that the short 10 amino acid linker used to fuse LIS1 to CUL4B may hinder the HLH domain to approach DDB1 (Supp. Figure 2A), the sequence alignment of LIS1 N-terminus with the helices of different HLH domains of diverse confirmed DCAFs shows that the first helix in LIS1 (residues 2-20) contains the conserved signature residues required to mediate an interaction with DDB1 (Słabicki et al., 2020) (Figure 2D). These data suggest that LIS1 uses two distinct surfaces to bind DDB1. One engages the DDB1 BPA-BPC cleft via its first conserved helix, and a second binding site comprises a conserved REKE loop in the DDB1 BPA domain interacting with the WD40 domain of LIS1.

**Figure 2:**
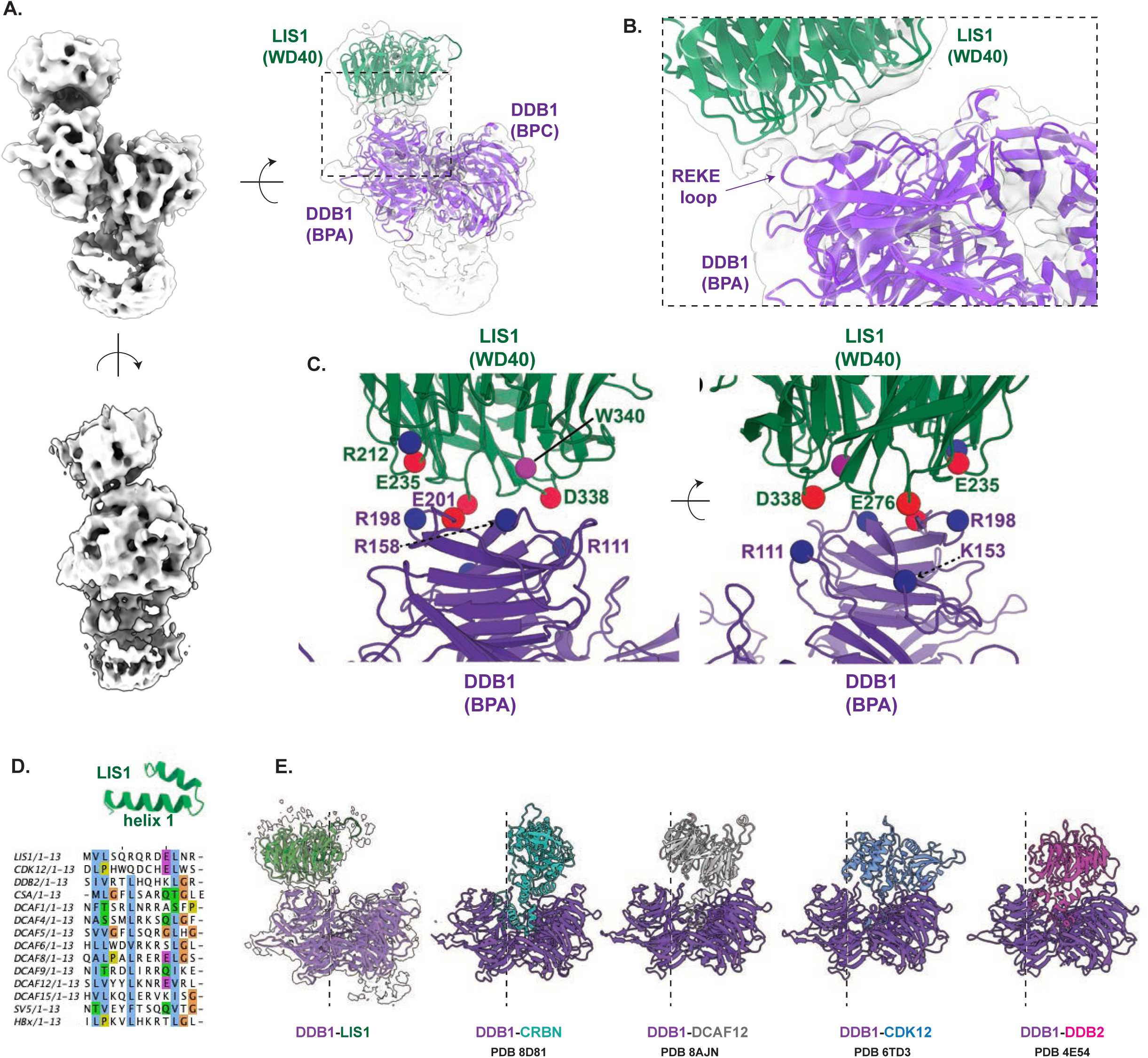
Identification of a unique REKE loop at the DDB1-LIS1 interface. **A.** Rotational views of the refined cryo-EM map at 4.3 Å-resolution. DDB1 (purple) and LIS1 (green) structures are fitted into the map as indicated. **B.** Zoom-in view of the LIS1-DDB1 interface, revealing a conserved REKE loop (arrow) located in the BPA domain of DDB1, which approaches the WD40 domain of LIS1. **C.** The interaction surface between LIS1 and the REKE loop in DDB1 contains charged residues. LIS1 (green) and DDB1 (purple) are shown in cartoon representation. Residues that are likely involved in the interaction are shown as spheres that are colored according to their side chain chemistry (negatively charged, red; positively charged, blue; aromatic, violet). **D.** Sequence alignment of reported HLH domains of different DCAFs. The HLH-fold in the amino terminal domain of LIS1 is indicated, highlighting helix 1 that interacts with CUL4B. **E.** Comparison of the described DDB1-LIS1 complex to atomic structures of canonical DDB1-DCAFs such as CRBN (cyan), DCAF12 (light gray), CDK12 (light blue), and DDB2 (magenta). The PDB number is specified underneath the corresponding complex. Note that in contrast to LIS1, known CUL4A DCAFs do not engage the DDB1 REKE loop.

Interestingly, comparing the DDB1-LIS1 structural model to atomic structures of conventional DCAFs revealed that the WD40 domain of LIS1 occupies a unique DDB1 interface, which is spatially removed from the commonly engaged BPC domain (Figure 2E). Indeed, even in DCAF structures such as DDB1-CDK12 and DDB1-DCAF12 that show WD40 or WD40-like domains to be spatially proximal to the BPA domain of DDB1, the REKE loop is not engaged. Therefore, we conclude that the conserved REKE loop may provide a previously unrecognized specificity interface preferentially occupied by CUL4B-specific DCAFs.

## The DDB1 REKE loop contributes to the DCAF preference of CUL4B

To establish that the REKE loop contributes to LIS1-DDB1 binding, we mutated this loop to either AAAA or EAAA, which based on structural analysis is predicted to disrupt LIS1 binding without altering DDB1 folding. As expected, the wild-type and DDB1 mutants express at equal levels and readily bind CUL4B-RBX1 (Supp. Figure 3A). Interestingly, decreased binding of LIS1 to DDB1 mutants was observed in (1) pulldown experiments after expressing the individual proteins in Hi5 insect cells (Supp. Figure 3B), and (2) using *in vitro* reconstitution of purified CRL4B complexes in the presence of wild-type DDB1 or the EAAA mutant with increasing concentrations of LIS1 (Supp. Figure 3C). In contrast, Alphafold multimer prediction of DDB1 (Δ-BPB) in complex with DCAF8 (iPTM 0.9) suggests that DCAF8 embeds its HLH in the DDB1 BPA-BPC cleft, while the WD40 domain interacts with the DDB1 BPC (Figure 3A) (Jumper et al., 2021). Overlaying this structure with the cryo-EM map of the DDB1-LIS1 complex confirmed that DCAF8 does not engage the DDB1 REKE loop at the BPA interface (Figure 3A).

**Figure 3:**
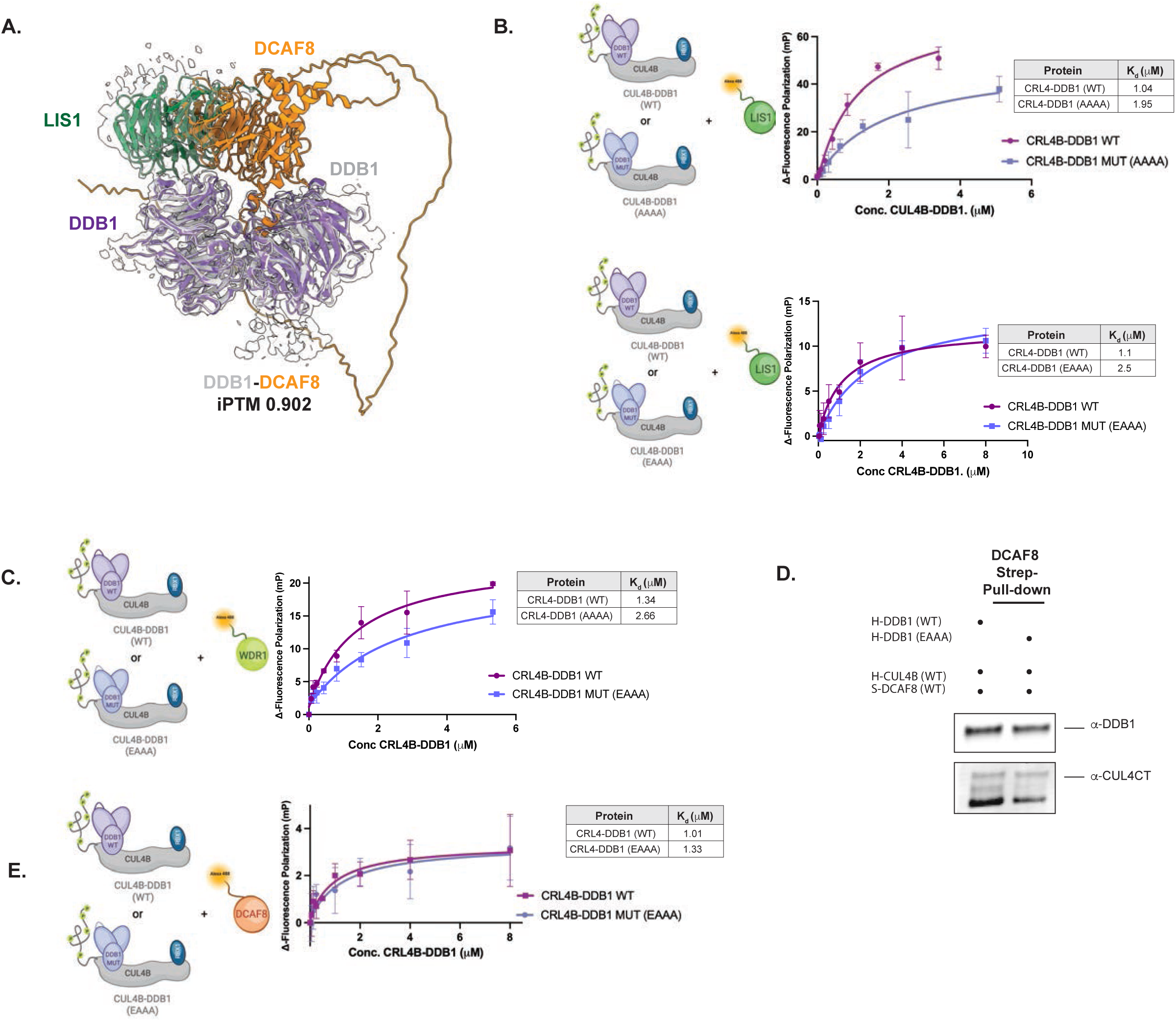
REKE loop mutations affect LIS1 binding but not the canonical substrate receptor, DCAF8. **A.** AlphaFold multimer prediction of DDB1 (gray) bound to DCAF8 (orange) (iPTM 0.92) overlaid on the DDB1 (purple)-LIS1 (green) cryo-EM structural model. Note that LIS1 and DCAF8 are predicted to use distinct binding sites. **B.** and **C**. Schematic diagram showing the experimental setup of the fluorescence polarization (FP) affinity assays (left). Wild-type (WT) DDB1 is shown in purple, the DDB1 REKE-mutants (AAAA or EAAA) in blue, LIS1 or WDR1 in green, and the Alexa488 fluorophore in glowing orange. FP binding curves were measured using increasing concentrations of CUL4B-DDB1 (WT; purple curves) and CUL4B-DDB1 mutant (AAAA or EAAA; blue curves) complexes in the presence of a fixed concentration of Alexa488-labeled LIS1 (**B**) or WDR1 (**C**). The quantified Kds (μM) are reported in the corresponding tables. **D.** Strep pull-down using extracts prepared from Hi5 insect cells expressing Strep-tagged DCAF8, His-tagged CUL4B/Rbx1, and His-tagged wild-type (WT) or the EAAA REKE mutant (MUT). **E.** Schematic diagram of the FP setup to measure affinities of CRL4B (WT vs. MUT) to DCAF8. Wild-type (WT) DDB1 is shown in purple, the DDB1 REKE-mutant in blue, DCAF8 in orange, and the Alexa488 fluorophore in glowing orange (left). FP binding curves were measured using increasing concentrations of CUL4B-DDB1 (WT; purple curve) or CUL4B-DDB1-EAAA (blue curve) complexes in the presence of a fixed concentration of Alexa488-labeled DCAF8. Quantified Kds (μM) are reported in the table.

To corroborate these data, we set-up quantitative fluorescence polarization (FP) binding assays using Alexa-488-labeled CUL4B-specific DCAFs LIS1 and WDR1, and the CUL4A-specific DCAF8 for control (Reichermeier et al., 2020).We first used FP assays to measure the interaction of wild-type CUL4B-RBX1-DDB1 (CRL4B DDB1 WT) and CUL4B-RBX1-DDB1 mutants AAAA or EAAA (CRL4B DDB1 mutant) in the presence of Alexa-488 labeled LIS1. Interestingly, these assays showed that mutationally disrupting the DDB1-REKE loop decreased the binding affinity of LIS1 approximately 2-fold (Figure 3B). Similarly, the Kd of Alexa-488 labeled WDR1 binding to CUL4-RBX1 complexes with wild-type DDB1 dropped almost 2-fold in the presence of the DDB1 EAAA REKE loop mutant (Figure 3C). In contrast, strep-pulldown assays with purified DCAF8 and CUL4B-RBX1 complexes containing wild-type DDB1 or the EAAA mutant showed no change in band intensity, implying comparable binding affinities (Figure 3D, Supp. Figure 3D). In agreement, Kd measurements using FP assays confirmed almost equal binding affinities of Alexa-488 labeled DCAF8 with wild-type CRL4B-DDB1 (WT) and CRL4B-DDB1 REKE mutant complexes (Figure 3E). The DDB1 REKE loop is highly conserved across various species, including unicellular organisms like *S. pombe* (Supp. Figure 3E), indicating its significance for DDB1 function. Taken together, these results indicate that CUL4B-specific DCAFs engage the novel DDB1 REKE loop, while CUL4A specific complexes bind DCAFs through a distinct binding surface located in the BPC domain of DDB1.

DDA1 also binds the BPA domain of DDB1 and bridges the interaction to a small subset of CUL4A DCAFs such as DCAF5, CSA, and DCAF15. Interestingly, in published atomic structures, residue R57 in DDA1 interacts with E201 in DDB1 across three CRL4 distinct complexes (Supp. Figure 3F and G) (PDB 6SJ7, PDB 8QH5, PDB 8TL6) (Burgess et al., 2025; Bussiere et al., 2020; Llerena Schiffmacher et al., 2024; Radko-Juettner et al., 2024), implying that DDA1 engages the REKE loop. Thus, our findings indicate that the REKE loop in DDB1 represents a novel interaction surface relevant for selective recruitment of either CUL4B-specific DCAFs or DCAFs that engage CUL4A via DDA1.

## The unique N-terminal CUL4B extension directly interacts with CUL4B-specific DCAFs

Surprisingly, LIS1 interacted with DDB1 alone with a low affinity compared to the entire CRL4B complex, and the absence of the CUL4B scaffold reduced LIS1 sensitivity to the DDB1 REKE loop mutation (Figure 4A). Structurally the DDB1 BPA domain is in close spatial proximity to the unique CUL4B N-terminus, suggesting that CUL4B may contribute to the DDB1-LIS1 interaction. To gain deeper insight into the interaction network, we performed cross-linking mass spectrometry analysis (XL-MS) of the CRL4B^LIS1^ complex (Figure 4B). As expected, many crosslinks were detected between the CUL4B scaffold and the RBX1 and DDB1 subunits. Importantly, extensive interactions were also exposed between CUL4B and LIS1, linking LIS1 to the C-terminus and the unique N-terminal CUL4B extension. In particular, Lys55 (K55) within the CUL4B N-terminus mediates several crosslinks to the WD40 domain of LIS1. This lysine is in a highly conserved region and in close proximity to the well-characterized proline 50 residue (Figure 4B, lower panel), which is commonly mutated in XLID patients. These data suggest that LIS1 directly contacts the N-terminal extension of the CUL4B scaffold. To test this notion, we performed FP binding assays with CUL4B-RBX1 alone, in absence of DDB1, and measured binding affinities to Alexa-488 labeled LIS1 or WDR1, and for control DCAF8. As expected, no interaction of CUL4B and DCAF8 was observed in the absence of DDB1 (Figure 4C), confirming that DCAF8 recruitment is fully dependent on DDB1. Strikingly, however, LIS1 readily interacted with CUL4B-RBX1, and an even stronger Kd of approx. 0.7 μM was observed for WDR1 and CUL4B-RBX1 (Figure 4C). Taken together, we conclude that recruitment of LIS1 and WDR1 into the CRL4B complex is mediated by direct binding to the N-terminal CUL4B extension and the REKE loop of DDB1.

**Figure 4:**
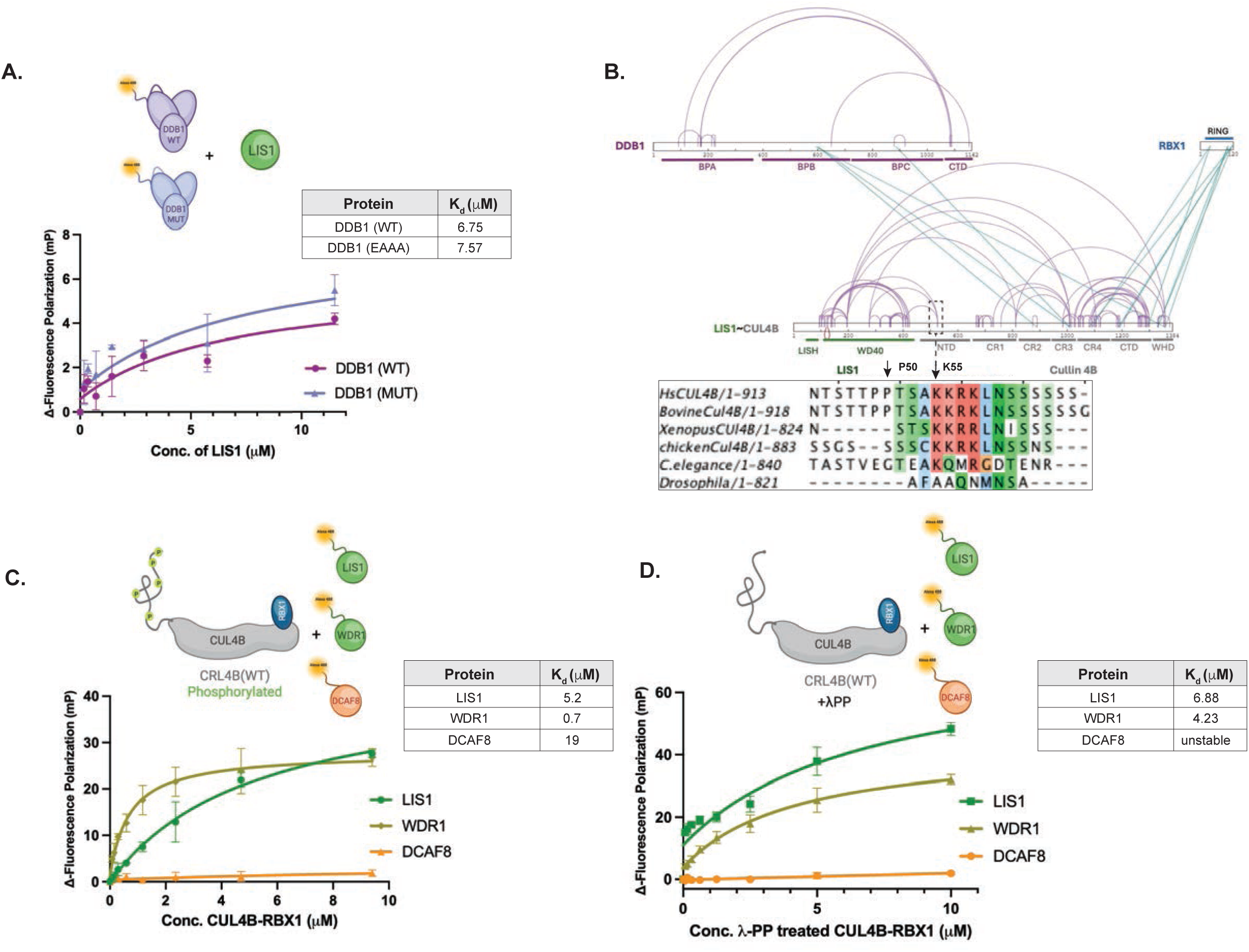
The CUL4B N-terminus directly interacts with the CUL4B-specific DCAFs, LIS1 and WDR1, but not with DCAF8. **A.** Schematic diagram of the Fluorescence polarization (FP) assays to measure binding affinities of DDB1 and LIS1. LIS1 is shown in green, wild-type (WT) DDB1 in purple, the DDB1 REKE-mutant (MUT) in blue, and the Alexa488 fluorophore in glowing orange (left). FP binding curves were measured using increasing concentrations of Alexa488-labeled DDB1 (WT; purple curve) or the DDB1-EAAA mutant (MUT; blue curve) in the presence of a fixed concentration of LIS1. Quantified Kds (μM) are listed in the table. **B.** Crosslinking-mass-spectrometry analysis of the recombinant CRL4B^LIS1^ protein complexes. Cross-links among different subunits are indicated by green lines, and cross-links within the same subunit by purple lines. The different subunits with domain boundaries are colored as follows: DDB1 in purple, LIS1 in green, CUL4B in gray, and Rbx1 in blue. The area highlighted with a rectangular box shows a prominent cross-link between Lys55 (K55) in CUL4B and the WD40 domain of LIS1. The conservation of the N-terminal KKRK motif in CUL4B from different species is displayed below, highlighting K55 and the P50 residue mutated in XLID patients. Visualized with xiNET (Combe et al., 2015). **C.** and **D**. Schematic diagram of FP binding assays measuring the affinity of phosphorylated or dephosphorylated (λPP treated) CUL4B-RBX1 complexes without DDB1 to Alexa488 (orange)-labeled LIS1, WDR1 (green) and for control, DCAF8 (red). FP binding curves were measured using increasing concentrations of phosphorylated (**C**) or dephosphorylated (**D**) CUL4B-RBX1 in the presence of a fixed concentration of LIS1 (green curve), WDR1 (asparagus green curve) or DCAF8 (orange curve). Quantified Kds (μM) are listed in the tables.

Interestingly, both LIS1 and WDR1 assemble CRL4B complexes specifically during mitosis, and this regulation is mediated by phosphorylation of multiple sites in the N-terminal extension of CUL4B (Stier et al., 2023). To test whether CUL4B phosphorylation directly regulates LIS1 and WDR1 binding, we treated purified CUL4B protein with λ-phosphatase, which efficiently removed phosphates (Supp. Figure 4A and B). While LIS1 binding was only slightly decreased upon dephosphorylation, the affinity of WDR1 dropped almost 5-fold (Figure 4C and D). As expected, no effect of CUL4B phosphorylation was observed for DCAF8. Moreover, CUL4B phosphorylation did not affect its interaction with DDB1, as we measured equal binding affinities of DDB1 to phosphorylated or dephosphorylated CUL4B-RBX1 (Supp. Figure 4C). Collectively, these results demonstrate that phosphorylation of the CUL4B N-terminal extension enhances binding of mitotic DCAFs.

### BRWD1 behaves structurally and biochemically like CUL4B-specific DCAFs

To systematically identify CUL4B-specific DCAFs, we took advantage of Hela Kyoto cell lines endogenously expressing HA-tagged CUL4B full length (FL) or truncated CUL4B (ΔN) and used MS-analysis to compare cellular proteins that interact with different affinity with wild-type CUL4B and CUL4B-ΔN lacking the unique N-terminal extension (Figure 5A). Each immunopurification was controlled for total protein abundance and the presence of comparable amounts of the HA-CUL4B bait (Supp. Figure 5A and B). We initially identified CUL4B interactors by comparing the co-immunoprecipitates of CUL4B and CUL4B-ΔN with the control purification, which yielded 47 specific interactors (Supp. Figure 5C and D, Supp. Table 1). Subsequently, the abundance of the interactors was normalized to the expression of the bait (HA-CUL4B-FL and HA-CUL4B- ΔN) and compared to evaluate the role of the N-terminal region to interactor recruitment. (Figure 5A). While this analysis identified few components that preferentially interact with truncated CUL4B, most known CRL regulators involved in CUL4 neddylation such as subunits of the COP9/signalosome were unchanged or slightly reduced in the absence of the N-terminal domain. Importantly, the comparison detected several DCAFs that bind CUL4B by a mechanism dependent on its N-terminal extension (Figure 5A, right panel). Many of these DCAFs were previously identified in a targeted approach as CUL4B-specific DCAFs (Reichermeier et al., 2020), except that our study missed AMBRA1 and BRWD3, as their binding was not affected by the CUL4B N-terminal deletion. Taken together, these data suggest that CUL4B-specific DCAFs may generally require direct binding to the N-terminal extension, despite the presence of a canonical HLH-motif.

**Figure 5:**
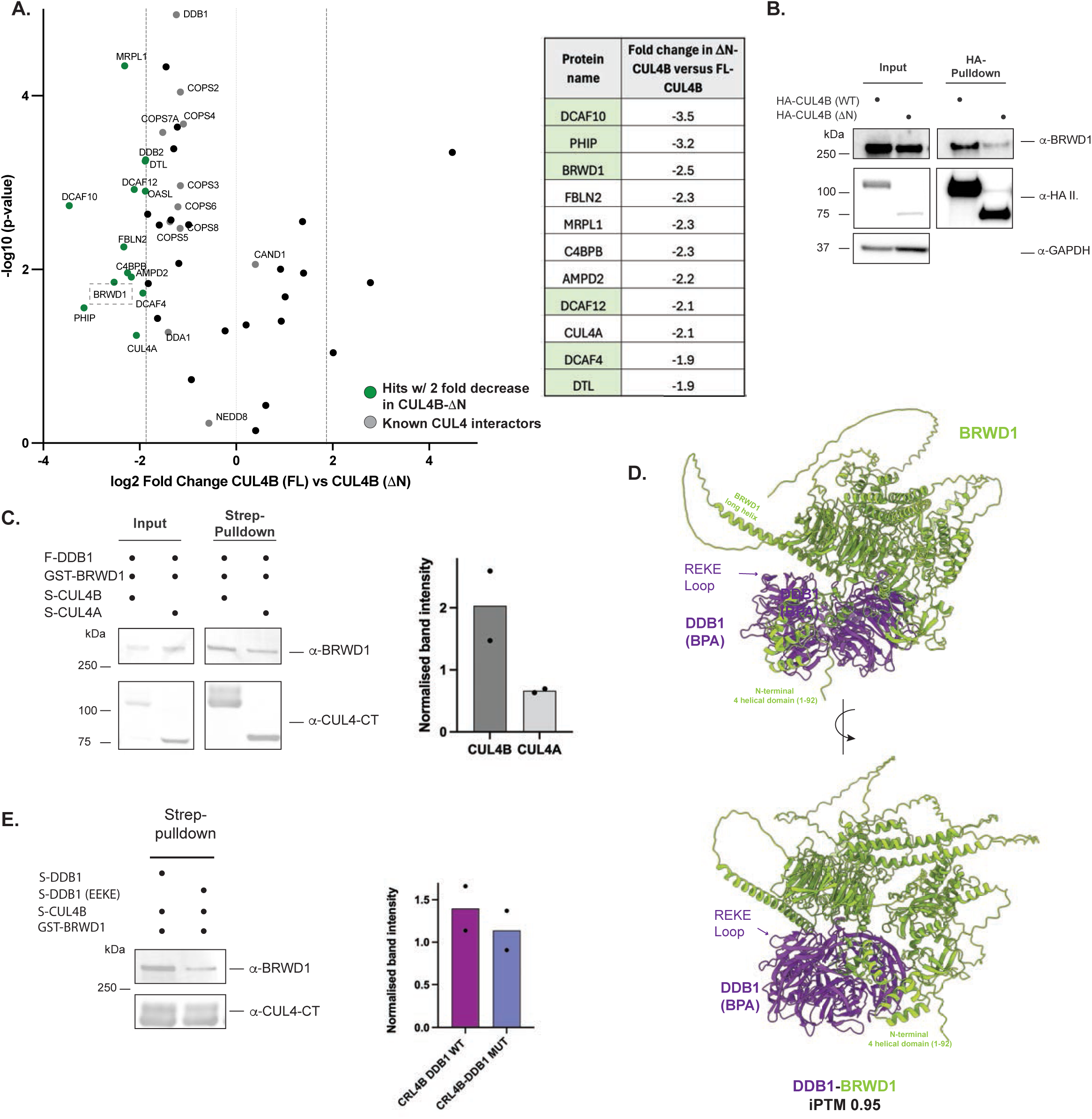
Identification of CUL4B-specific DCAFs including BRWD1 that require interaction with the CUL4B N-terminal extension. **A.** AP-MS pulldown experiment using extracts prepared from HeLa Kyoto cell lines stably expressing either HA-tagged full-length CUL4B (FL) or the truncated CUL4B mutant lacking its unique N-terminal extension (ΔN). Differential abundance of the 47 CUL4B interactors is reported as the log2 ratio between the ΔN and FL conditions. Proteins previously identified as CUL4B-specific DCAFs are shown in green (Reichermeier et al., 2020). Proteins levels are normalized to the bait abundance (CUL4B) and the significance is calculated from three biological independent replicates using a two-sided unpaired Student’s t’test. Proteins that significantly interact with the baits (p < 0.05) are indicated in bold, and BRWD1 proteins is highlighted in red. Error bars represent the standard error. **B.** Total extracts (input) prepared from HeLa Kyoto cell lines stably expressing either HA-CUL4B (FL) or HA-CUL4B (ΔN) were subjected to immunoprecipitation with HA antibodies (HA pulldown). The input fractions and immunoprecipitates were probed with antibodies against BRWD1, and for control HAII and GAPDH. **C.** Western blot of total extracts (input) and *in vitro* Strep-pull-down assays (Strep-pulldown) of Hi5 insect cells expressing Strep-tagged CUL4B or CUL4A together with FLAG-tagged DDB1 and GST-tagged BRWD1 (left). A secondary fluorescence antibody was used for detection with the LiCor Odyssey device. Fluorescent band intensities were quantified with Fiji from two biological replicates and displayed using prism software (right). Note that BRWD1 preferentially binds CUL4B compared to CUL4A. **D.** AlphaFold 3 structural prediction of DDB1 (Δ-BPB) bound to BRWD1 (Δ-1451-2182). iPTM 0.95. BRWD1 uses an extended helix that approaches the DDB1 REKE loop in the BPA domain. **E.** Western blot of *in vitro* pull-down assays of Strep-tagged CUL4B/RBX1 co-expressed in Hi5 insect cells with GST-tagged BRWD1 and either wild-type DDB1 (WT) or the indicated REKE loop mutant. A secondary fluorescence antibody was used for detection with the LiCor Odyssey device. Fluorescent band intensities are quantified using Fiji (right) between two biological replicates and displayed in prism software.

To corroborate these data, we focused further analysis on BRWD1, a large 260 kDa DCAF involved in chromatin remodeling and transcriptional regulation. Indeed, HA-immunoprecipitation experiments followed by immunoblotting validated that BRWD1 preferentially binds HA-CUL4B-FL compared to HA-CUL4B-ΔN (Figure 5B). We next overexpressed the subunits of CRL4B^BRWD1^ (CUL4B-RBX1-DDB1-BRWD1) and CRL4A^BRWD1^ (CUL4A-RBX1-DDB1-BRWD1) complexes in Hi5 insect cells and pulled on the strep-tagged CUL4 scaffold. The bound fractions were then immunoblotted for the presence of BRWD1 using a fluorescently-tagged secondary antibody and quantified with a LiCOR system (Figure 5C, left panel). Interestingly, BRWD1 incorporated much less into a complex with CUL4A compared to CUL4B (Figure 5C, right panel), consistent with the observed binding preference *in vivo*. A high-confidence AlphaFold 3 prediction of DDB1(Δ-BPB)-BRWD1(Δ1451-2182) (iPTM 0.95) identified a long helix on BRWD1 (748-796) approaching the BPA of DDB1 (Figure 5D) (Abramson et al., 2024). Indeed, residues within this long helix, such as R775 and E760 (Supp. Figure 5D) seem to interact with the REKE loop in DDB1, such as R111, R158 and E201 (Figure 2C). In addition, BRWD1 has an additional N-terminal α-helical domain composed of four short helices (1-92) that may further interact with the BPA domain of DDB1 (Figure 5D). Therefore, we tested whether the REKE loop mutations of DDB1 affect BRWD1 incorporation using pulldown assays with strep-tagged CRL4B complex. Consistently, BRWD1 incorporation into CRL4B complexes was reduced in the presence of DDB1-REKE mutants (Figure 5E), confirming that this surface contributes to BRWD1 binding. Similar results were obtained using a different strep-pulldown set-up, in which we immobilized purified CRL4B complexes on strep beads, and incubated the CRL4B-loaded beads *in vitro* with Hi5 lysates of cells overexpressing BRWD1. After washing, bound BRWD1 was then quantified by immunoblotting using a fluorescently-tagged secondary antibody (Supp. Figure 5D). Together, these data suggest that BRWD1 functions as a CUL4B-specific DCAF, stabilized by binding to the amino-terminal CUL4B domain and the REKE loop of DDB1.

## Discussion

Here we address why vertebrates express two CUL4 paralogs, e.g. CUL4A and CUL4B. We found that CRL4B employs a dual mechanism to preferentially bind specific DCAFs, namely its unique CUL4B N-terminal extension and the conserved REKE loop in DDB1. These novel interactions cooperate to different extent with the canonical HLH motif, which may be compensated however by phosphorylation of the CUL4B N-terminus. These results raise the possibility that additional CUL4B-specific DCAF substrate receptors may exist.

## CUL4B binds specific substrate receptors through a novel DDB1 interaction surface

Structural and biochemical analysis has shown that conventional DCAFs do not directly interact with the CUL4 scaffold but exclusively bind DDB1 through a conserved HLH or HLH-like domain. Indeed, this conserved structure inserts into a cleft formed by the BPA and BPC β-propeller domains of DDB1, thus providing specificity to assemble CUL4A and CUL4B complexes. These DCAFs show high affinity to bind DDB1 and are only soluble when complexed with DDB1. The CUL4B substrate receptor LIS1 contains conserved residues encompassing a canonical HLH motif for interaction with DDB1 (Figure 2D). The first helix overlaps with the LISH motif, which is known to dimerize. However, in the CRL4B^LIS1^ structure, LIS1 exists as a monomer, and when interacting with DDB1, the LISH motif is not accessible for dimerization. This mutually exclusive binding mechanism may explain how LIS1 can adopt different functions, on the one hand regulating dynactin at the cell cortex, and on the other hand assembling into a CRL4B^LIS1^ E3 ligase complex regulating spindle positioning during mitosis.

Surprisingly, we found that CUL4B-specific substrate receptors engage a previously unrecognized REKE loop located at the surface of the DDB1 BPA β-propeller. While the HLH or HLH-like domains remain a primary interaction interface with DDB1, an intact REKE loop contributes to the recruitment of CUL4B- but not CUL4A-specific DCAFs such as DCAF8. Consistent with its functional relevance, the REKE loop is highly conserved among eukaryotes (Supp. Figure 3E), implying that this alternative DDB1 binding mechanism is widespread. CUL4B specific DCAFs may employ different motifs to interact with the REKE loop of DDB1. For example, LIS1 engages charged residues located in its WD40 domain, while in BRWD1 residues located within a long helix direct DDB1 binding. CUL4B specific DCAFs may also differ to which extent they engage the REKE loop in addition to the canonical HLH-motif and direct binding to the N-terminal extension.

Interestingly, the REKE loop is not accessible for DCAF binding in the absence of CUL4B, suggesting that DDB1 undergoes a conformational change upon CUL4B binding that favors recruitment of specific substrate receptors. This conformation seems absent when DDB1 is incorporated in CRL4A complexes (Figure 6), and we thus speculate that this unique DDB1 conformation is enforced by the presence of the unique CUL4B N-terminus, which is near the BPA domain. The REKE loop may thus act as a molecular switch to facilitate CUL4B-specific DCAFs recruitment.

**Figure 6:**
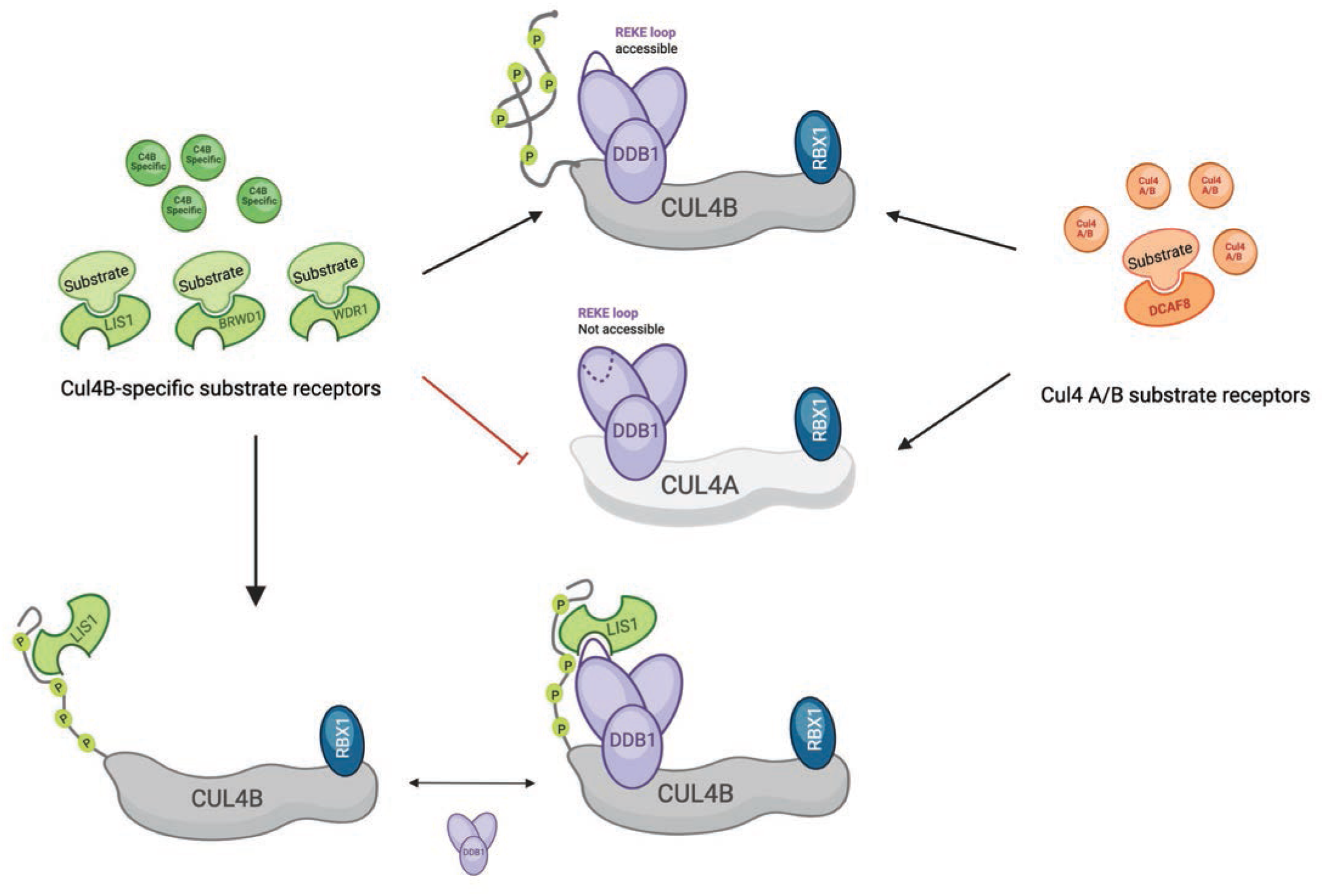
Model depicting specificity determinants to recruit CUL4B specific DCAFs The REKE loop in the BPA domain of DDB1 (violet) provides a novel interaction surface, which is accessible when DDB1 is bound to CUL4B/Rbx1 (pointed arrow) but not CUL4A/Rbx1 (blunt arrow). This conformational switch is most likely triggered by the N-terminal extension of CUL4B. In addition, CUL4B specific substrate receptors (green) such as LIS1 directly bind the unique CUL4B N-terminus, and this interaction can be regulated by phosphorylation. Together, these mechanisms ensure specificity of CUL4A and CUL4B substrate receptors (orange and green, respectively).

Of note, the functional relevance of the REKE-loop may not be limited to CUL4B-specific DCAFs. Indeed, binding of some DCAFs to DDB1 is dependent on DDA1, and the crystal structures of DDA1-DDB1-CRL4A complexes with DCAF15 (PDB: 6SJ7), CSA (PDB: 8QH5) and DCAF5 (PDB: 8TL6) revealed that residues 53-61 adopt a helix, in which arginine 57 of DDA1 is as close as 2 Å to the glutamate residues of the REKE loop of DDB1 (Supp. Figure 3. F and G). This implies that like CUL4B specific DCAFs, DDA1 may exploit the REKE loop to enhance recruitment of some DCAFs into CUL4A complexes.

## CUL4B directly binds specific substrate receptors through its unique N-terminal extension

Strikingly, we found that the amino-terminal extension of CUL4B directly interacts with multiple CUL4B-specific substrate receptors. Indeed, WDR1 can bind CUL4B with an affinity of ∼0.7 mM and LIS1 with ∼6 mM, and it may thus require stronger interaction interfaces with the CUL4B scaffold. In contrast to LIS1, WDR1 lacks a canonical HLH motif, as AlphaFold predictions of the WDR1-DDB1 complex produced only unreliably low iPTM scores (0.2-0.3), and a recent cryo-EM structure of WDR1 in the cofilactin complex did not identify a HLH fold (Oosterheert et al., 2025). Increased binding of WDR1 to the N-terminal CUL4B domain may thus compensate for the lack of the HLH-mediated interface with DDB1. As expected, the CUL4A-specific DCAF8 does not interact with CUL4B, consistent with previous data that its CRL4 recruitment is solely mediated by DDB1. Due to the limited resolution of the CRL4^LIS1^ cryo-EM model, we could not resolve the atomic structure of this dynamic low-complexity domain. However, crosslinking-MS experiments confirmed that K55 must be near the WD40 domain of LIS1. K55 is also close to T49, which regulates phosphorylation-dependent recruitment of LIS1 during mitosis. Likewise, mitotic phosphorylation of WDR1 is critical to enhance its ability to bind CUL4B. In contrast, BRWD1 binds CUL4B during cell cycle stages other than mitosis, suggesting that phosphorylation of the N-terminus may provide an additional mechanism to modulate specificity of CRL4B complexes. Indeed, the N-terminal domain encompasses multiple phosphorylation residues, and only some of them are upregulated during mitosis, raising the possibility that a CUL4B phosphorylation code may contribute to select substrate-specific receptors in space and time.

## Unbiased identification of CUL4B specific substrate receptors

Interestingly, our unbiased MS-analysis comparing proteins predominantly interacting with CUL4B in the presence of its N-terminus, detected a set of DCAFs strongly overlapping with a targeted study probing a set of previously known DCAFs for their preference to bind CUL4A or CUL4B (Reichermeier et al., 2020). This striking similarity supports the conclusion that the unique CUL4B N-terminus provides specificity in recruiting specific substrate receptors. Among them, BRWD1/DCAF19 emerged as a common hit, and it will thus be interesting to explore its CUL4B-specific functions. Available data suggest that BRWD1 is a chromatin-associated substrate-receptor, which based on its two BROMO-domains is predicted to regulate chromatin remodeling, and may thus link histone acetylation and CUL4B-dependent ubiquitination. Surprisingly, we did not detect AMBRA1 or BRWD3 as CUL4B-specific DCAFs, as their binding was not affected by the CUL4B N-terminal deletion. It is thus possible that additional specificity determinants exist that may contribute to their CUL4B preference.

Our unbiased MS-approach also has the potential to identify proteins which have not previously been recognized as DCAFs. Along these lines, we detected the 2’’-5’’-Oligoadenylate Synthetase Like (OASL) protein, involved in viral infections, and the C4b-binding protein (C4BP), implicated in degrading glycosylated proteins in apoptotic and necrotic cells. Further work will be required to examine whether these proteins indeed function as CUL4B-specific DCAFs. Likewise, we did not find the mitotic CUL4B substrate-receptors LIS1 and WDR1, most likely because mitotic cells are underrepresented in unsynchronized cultures. Thus, analyzing cell extracts prepared from cells arrested at different cell cycle stages or exposed to specific environmental conditions may be warranted to detect novel CUL4B-specific DCAFs that may be regulated in response to intra- or extracellular signals.

## Materials and Methods

### Antibodies and reagents

For western blot analysis following primary antibodies were used: anti-LIS1 (mouse, Santa Cruz Biotechnology); anti-CUL4-CT (rabbit) (Olma et al., 2009); anti-CUL4B (rabbit, Sigma); anti-DDB1 (mouse, BD Biosciences); anti-GAPDH (mouse, Sigma); anti-HA.II (mouse or rabbit, both Covance); anti-BRWD1 (Rabbit, Sigma); anti-FLAG (mouse, Sigma). For western blot anti-mouse or rabbit IgG-HRP (both Biorad) and for fluorescent LiCor scans, fluorescent secondary antibodies IRDye® 800CW Goat anti-Rabbit and IRDye® 680LT Goat anti-Mouse (LiCorbio) were used.

### Protein expression and purification

The fusion CUL4B-LIS1 gene was designed and then split into several gene fragments, which were ordered (IDT) and assembled by Gibson assembly (Gibson et al., 2009). DDB1 mutants were cloned by Gibson assembly of the mutated gene fragment into the DDB1 vector. Recombinant baculoviruses were prepared in *Spodoptera frugiperda* (*sf9*) cells using the Bac-to-Bac system (Life Technologies). Recombinant protein complexes were expressed in *Trichoplusia ni* High Five cells by co-infection of single baculoviruses. Protein complexes were expressed by co-infection of Hi5 cultures with 17.5 ml per liter of the corresponding baculoviral preparation. The CRL4B^LIS1^ preparation was expressed using Strep-tagged CUL4B-LIS1, Flag-tagged DDB1, and His-tagged RBX1 baculoviruses. CRL4B complexes with wild-type (WT) or mutated DDB1 were expressed with N-terminal Strep (II) tagged DDB1 and CUL4B and His-tagged RBX1. CUL4B-RBX1 preps were expressed using baculoviruses of Strep (II) tagged CUL4B and His-tagged RBX1. Full length WDR1, LIS1, or DCAF8, as well as DDB1 (WT) or the DDB1 EAAA mutant were expressed separately, each with an N-terminal Strep (II) tag, or Strep II-Avi tag. Cultures were incubated shaking at 27 °C for 1.5 days. Every liter was lysed with 30 mL of lysis buffer (Tris pH 8.0 50 mM, NaCl 300 mM, EDTA 0.5 mM, tris(2-carboxyethyl)phosphine (TCEP) 0.5 mM, Triton X-100 1%, protease inhibitor cocktail (Roche Applied Science) and 1 mM phenylmethanesulfonyl fluoride PMSF, and glycerol 10%. After lysis 5 μl benzonase (NEB) were added and stirred for 15 mins at room temperature (RT). Sonication with 50% amplitude was performed for 9 rounds of 30 seconds each with 1.5 seconds pulse and 1 second off and 2 mins break on ice between each two rounds using Bandelin Sonopuls sonicator. Lysates were cleared by ultracentrifugation for 45 min at 40,000 *g*. The supernatant was loaded on equilibrated Strep-Tactin (IBA life sciences). The CRL4B-LIS1 fusion preparation was treated with 10 mL of bioblock (IBA life sciences) and incubated for 25 mins to get rid of a biotinylated-contaminant. Several washes (>10 CV) with wash buffer (Tris pH 7.5 50 mM, NaCl 300 mM, TCEP 0.5 mM, and glycerol 10%) were performed and the fusion protein was eluted in elution buffer (Tris pH 8.0 50 mM, NaCl 300 mM, TCEP 0.5 mM, glycerol 10%, and desthiobiotin 5 mM). Elution fractions were further purified via ion exchange chromatography (Poros HQ 50 µm, Life Technologies) and subjected to size-exclusion chromatography in a buffer containing 50 mM HEPES pH 7.4, 200 mM NaCl, 0.5 mM TCEP, and 2-10% glycerol. Pure fractions, as judged by SDS-PAGE, were collected, concentrated using 10,000 MW cut-off centrifugal devices (Amicon Ultra) and stored at -80 °C.

### *In vitro* auto-ubiquitination assays

*In vitro* auto-ubiquitination assays were performed by mixing 1 μM CRL4B-LIS1 complex with a reaction mixture containing 0.2 μM E1 (UBA1, BostonBiochem), 1 μM E2 (UBCH5a, BostonBiochem) and 50 μM ubiquitin (Ubiquitin, BostonBiochem). Reactions were carried out in 50 mM HEPES pH 7.4, 200 mM NaCl, 10 mM MgCl_2_, 0.2 mM CaCl_2_, 3 mM ATP, 0.5 mM TCEP, and incubated for 90 min at 32 °C. Reactions were stopped with SDS loading dye and analyzed by western blot using anti-LIS1 antibodies.

### CryoEM grid preparation

In order to increase the stability of the CRL4B^LIS1^ complex, a gradient fixation (GraFix) protocol was applied (Stark, 2010). A glycerol gradient (10%-40% w/v) in the presence of the cross-linker glutaraldehyde (0.25% v/v) was prepared using a Gradient Mixer machine, 150 μL buffer cushion of 7% glycerol was added on the top, and left for 30 minutes to settle at 4°C. 100 μL of 9 mg/ml of purified CRL4B^LIS1^ was loaded on the gradient, followed by ultracentrifugation (SW40Ti rotor) at 37,000 xg for 18 h at 4 °C. Peak fractions containing CRL4^LIS1^ were collected and cross-linking was quenched by adding 0.1M glycine at pH 7.5. glycerol removal and buffer exchange was performed by Zeba spin column equilibrated with buffer containing 50 mM HEPES pH7.4, 200 mM NaCl, 0.5 mM TCEP, and 0.05% NP40. 4 μL sample (0.15-0.3 mg/ml) was applied on glow discharged R1.2/1.3 and R2/2 Quantifoil holey grids manually floated with a 1.1 nm monocarbon layer (Quantifoil Micro Tools GmbH, Grosslöbichau, Germany).

### In vitro pull-down assays

For pull-down assays in Hi5 cells, 150 μl of corresponding baculoviruses per 10 ml of Hi5 cells were used. Cultures were co-infected with baculoviruses as follows: (1) His-DDB1 (WT) or His-DDB1 mutant (AAAA), Strep-DCAF8, and His-CUL4B, (2) His-DDB1 (WT) or His-DDB1 mutant (AAAA) together with Strep-DCAF8 in the absence of CUL4B, (3) FLAG-DDB1, GST-BRWD1, Strep-CUL4B or Strep-CUL4A, (4) GST-BRWD1, Strep-CUL4B with Strep-DDB1 (WT) or Strep-DDB1 mutants (EEKE or EAAA) (5) FLAG-LIS1, Strep-CUL4B with Strep-DDB1 (WT) or Strep-DDB1 mutants (AAAA or EAAA). Infected cells were incubated at 27 °C for 48 h and lysed by sonication in a buffer containing Tris-HCl pH 8, 200 mM NaCl and 0.5 mM TCEP, including 0.1% Triton X-100, 1x protease inhibitor cocktail (Roche Applied Science), 1 mM PMSF, and 10% glycerol. Lysates were cleared by centrifugation at 14,000 *g* for 30 min. Total protein concentration was normalized to the lowest concentration and 1 ml of soluble protein fractions were incubated for 1 h at 4 °C with 20-30 μl Strep-Tactin Macroprep beads (IBA life sciences) or FLAG beads (Sigma). Beads were washed three times with lysis buffer, bound proteins eluted in 20-30 μl of SDS loading dye and heated at 80 °C for 10 min.

For pull-down assays using purified protein complexes, the Strep-tag was cleaved from LIS1 (WT) using His-tagged TEV-protease. TEV was then removed by loading the cleaved sample on Ni-NTA beads (Sigma). Strep-tagged CUL4B-RBX1 at 0.25 μM was mixed with 0.25 μM Strep-tagged DDB1 (WT or EAAA mutant) and incubated for 10 min on ice. Increasing concentrations of LIS1 were added at 0.2, 0.8, 3 and 5 μM. The reactions were loaded on equilibrated 10 μl magnetic Strep-resin (IBA life science) and mixed by shaking at room temperature. Beads were washed 3 times with 500 μl of buffer containing HEPES pH 7.2 50 mM, NaCl 200 mM, and TCEP 0.5 mM, followed by the addition of 7.5 μl SDS loading dye and incubation for 10 min at 80 °C. Reactions were loaded on 4–15% Criterion™ TGX Stain-Free™ Protein Gel, 26 well (BioRad). Transfer was done using the Trans-blot Turbo system (BioRad).

For pull-down assays with purified protein complexes with BRWD1, 150 mL of Hi5 cells were infected with 1.5 mL of GST-BRWD1 baculovirus. Cultures grew for two days at 27 °C and were harvested by centrifugation at 1500 x g for 10 min. The pellet was lysed in 8 mL lysis buffer (Tris pH 7.5, NaCl 300 mM, TCEP 0.5 mM, 5% glycerol, PMSF, Protease inhibitor cocktail) followed by sonication for 6 rounds, with 1.5 s pulse for 20 seconds per round. After centrifugation at 21,000 xg for 45 mins, the reaction was split in two falcon tubes with 4.5 mL of lysate in each. A stock of 5.3 μM of purified Strep-tagged CUL4B-RBX1-DDB1 (WT) and Strep-tagged CUL4B-RBX1-DDB1 (EAAA) was prepared and incubated with equilibrated 100 μL Strep Macro beads (IBA life sciences) for 15 minutes. The CRL4-loaded beads were transferred to the 4.5 mL of lysate and incubated for 2 h at 4 °C. Beads were washed with 10 column volumes, bound fractions were eluted with SDS loading dye and analyzed by western blot.

### λ-Phosphatase treatment

4.2 μM of CUL4B-RBX1 was incubated in 1x PMP buffer supplemented with 1 mM MnCl_2_, and 1200 U λ-Phosphatase (NEB) for 2 h at room temperature.

### Biotinylation

Purified StrepII-Avi-tagged LIS1, StrepII-Avi-tagged DCAF8, StrepII-Avi-tagged DDB1 (WT) or StrepII-Avi-tagged DDB1 mutant (EAAA) were biotinylated *in vitro* at an amount of 9 mg for LIS1, 1.7 mg for DCAF8, and 3 mg for DDB1. BirA was added to a final concentration of 1/20 of the protein concentration to be biotinylated, with 0.2 mM D-Biotin in 50 mM HEPES pH 7.4, 200 mM NaCl, 10 mM MgCl_2_, 0.5 mM TCEP and 20 mM ATP. The reaction was incubated for 3 h at room temperature and at 4 °C overnight. Biotinylated proteins were purified by size exclusion chromatography and stored at −80 °C. Biotinylated WDR1 was used as previously described (Stier et al., 2023). Biotinylated proteins were mixed at 1:1 molar ratio with Streptavidin, Alexa Fluor™ 488 conjugate (Thermo Fischer) followed by buffer exchange using Zeba Spin columns equilibrated with Assay Buffer (HEPES pH 7.4 50 mM, NaCl 200 mM, D-Trehalose 25 mM, Triton X 0.01%, BSA 0.1 mg/ml and 0.5 mM TCEP).

### Fluorescence Polarization

26 nM of Alexa-labeled LIS1, WDR1, DCAF8, DDB1 (WT), or DDB1 (EAAA) were titrated with increasing concentrations of CUL4B-RBX1-DDB1 (WT), CUL4B-RBX1-DDB1 mutants (EAAA or AAAA), LIS1 (WT), and phosphorylated or dephosphorylated CUL4B-RBX1, as indicated. Reactions were incubated at room temperature in 384-well plates in a BMG Clariostar plate reader (BMG LabTech). Binding affinities were calculated by measuring the change in fluorescence polarization. Data were plotted and analyzed using GraphPad Prism assuming a single site binding saturation (Y=Bmax*X/(Kd + X)+C).

## Cell culture, immunoprecipitation and western blot experiments

HeLa Kyoto cells stably endogenously expressing HA-CUL4B (FL) or HA-CUL4B (ΔN) were prepared as described previously (Stier et al., 2023). Cells were grown in NUNC cell culture dishes in Dulbecco’s modified medium (DMEM) from Invitrogen supplemented with 10% FBS and 1% Penicillin-Streptomycin-Glutamine 100x (PSG, Life Technologies). Cells were harvested ∼48h after splitting and lysed in buffer containing 40 mM Tris HCl pH 7.4, 120 mM NaCl, 1mM EDTA, 0.3% CHAPS, 0.5 mM TCEP, 1x PhosSTOP, and 1x Complete Protease Inhibitor Cocktail (Roche), 1 mM PMSF, and 10% glycerol. Lysates were cleared by centrifugation for 30 min at full speed and protein concentrations were normalized to lowest protein concentration with lysis buffer. For immunoprecipitation experiments, lysates were loaded on HA-beads and incubated for 1 h at 4 °C. Beads were then washed three times with the lysis buffer, and bound proteins eluted with SDS loading dye.

For immunoblot analysis, bound proteins were eluted with SDS loading dye and incubated 10 mins at 80 °C. Samples were resolved by standard SDS-PAGE or NuPAGE 4-12% Bis-Tris Protein Gels (Invitrogen) before transfer onto Immobilon-PVDF or Nitrocellulose transfer membranes (Millipore). Before incubation with the respective primary antibodies, membranes were blocked in 5% milk-PBST (MIGROS) for 1h, followed by primary antibodies incubation overnight at 4 °C. Secondary antibodies were anti-mouse IgG HRP (170-6516, Bio-Rad) or goat anti-rabbit IgG HRP (170-6515, Bio-Rad). Proteins were visualized with SuperSignal™ West EC Chemiluminescent Substrate solution (Thermo Fisher) and scanned on a Fusion FX7 imaging system (Witec AG). For re-probing, blots were stripped in ReBlot Plus stripping buffer (2504 Millipore) and washed several times in PBST.

For AP-MS analysis, proteins were digested after purification using Filter Assisted Spin Proteolysis (FASP) as preciously described (Uliana et al., 2023; Wiśniewski et al., 2009). Briefly, proteins were loaded onto 10 kDa MWCO spin columns (Vivacon 500, Sartorius) and centrifuged at 9,000 × g until dry. Subsequently. samples were resuspended in 8 M urea, reduced with 5 mM TCEP, and alkylated with 10 mM IAA, followed by two washing with 0.1 M ammonium bicarbonate. Proteins were digested with 0.4 μg LysC (Wako) for 3 hours and subsequently with 1.2 μg trypsin (Promega) overnight. Proteolysis was quenched by addition of 5% formic acid and peptides collected by centrifugation (8,000 × g, 20 min). Prior to MS analysis, peptides were purified using C18 cleanup columns (7–70 μg capacity; The Nest Group) according to the manufacturer’s instructions, dried, and resuspended in 2% acetonitrile containing 0.1% formic acid.

## Mass Spectrometry data acquisition

For AP-MS experiment, LC-MS/MS analysis was performed on an Orbitrap QExactive+ mass spectrometer (Thermo Fisher) coupled to n EASY-nLC-1000 liquid chromatography system (Thermo Fisher). Peptides were separated using a reverse phase column (75 μm ID x 400 mm New Objective, in-house packed with ReproSil Gold 120 C18, 1.9 μm, Dr. Maisch GmbH) across a gradient from 3% to 25% in 110 minutes and from 25% to 40% in 10 minutes (buffer A: 0.1% (v/v) formic acid; buffer B: 0.1% (v/v) formic acid, 95% (v/v) acetonitrile). The DDA data acquisition mode was set to perform one MS1 scan followed by a maximum of 20 scans for the top 20 most intense peptides with MS1 scans (350-1500 m/z, R=70’000 at 400 m/z, AGC target = 3e6 and max. IT = 64ms), HCD fragmentation (NCE=25%), isolation windows (1.4 m/z) and MS2 scans (R=17’500 at 400 m/z, AGC = 1e5 and max. IT= 55ms). A dynamic exclusion of 30s was applied and charge states lower than two and higher than seven were rejected for the isolation.

## Data analysis of AP-MS experiment

Acquired data were searched using the MaxQuant software package version 1.5.2.8 embedded with the Andromeda search engine (Cox & Mann, 2008) against the Homo Sapiens 9606 reference dataset (http://www.uniprot.org/, downloaded on 06.04.2021, 20’388 proteins) extended with reverse decoy sequences. The search parameters were set to include only full tryptic peptides, maximum two missed cleavage, carbamidomethylation of C as static peptide modification, oxidation (M) as variable modification (maximum three variable modifications per peptide). The “match between runs” feature was enabled. The MS and MS/MS mass tolerance was set to 10 ppm, and FDR was controlled at 1% at PSM and protein level. Protein abundance was determined by label free quantification (LFQ) from the intensity of at least two peptides. Intensity values were median normalized and only proteins identified in triplicate in at least one condition were considered (in total 1476 proteins). Imputation of missing values (∼3%) was performed sampling the abundance from a normal distribution generated with 1% less intense values. Differential analysis of CUL4B and CUL4B ΔN against the control purification was performed to identify CUL4B interactors using as filtering criteria a log2FC> 1 and pvalue < 0.05 in at least one comparison (two-sided unpaired Student’s t-test, N= 3 biological replicates). These criteria led to the identification of 47 interactors. The abundance of CUL4B interactors was normalized to the abundance of the bait protein, and normalized values were compared between CUL4B and CUL4B ΔN conditions using a two-sided unpaired Student’s t-test (n = 3 biological replicates). The entire dataset, including raw data, generated tables and scripts used for the data analysis are available in the PRIDE repository (PXD074312)

## Cross-linking mass spectrometry

For the cross-linking experiments, the CRL4B^LIS1^ complex was prepared at an approximate protein concentration of 1 µg/µl in 50 mM HEPES, 200 mM NaCl, 0.5 mM TCEP, 10% glycerol. 50 or 150 µl were cross-linked at a final concentration of 0.25 mM disuccinimidyl suberate (DSS-d_0_/d1_2_, Creative Molecules) for 45 min at 25 °C with mild shaking (750 rpm). The reaction was quenched by addition of ammonium bicarbonate to a final concentration of 50 mM and incubation for additional 45 min at 25 °C. The samples were evaporated to dryness in a vacuum centrifuge and dissolved in 8 M urea for reduction and alkylation steps using tris(2-carboxyethyl)phosphine and iodoacetamide, respectively. Digestion was performed with endoproteinase Lys-C (Fujifilm/Wako, 2 h, 37 °C, 1:100) and trypsin (Promega, overnight, 37 °C, 1:50). Digests were brought to a pH <3 by addition of pure formic acid, purified by solid-phase extraction using 50 mg Sep-Pak tC18 cartridges (Waters) and fractionated by size-exclusion chromatography (SEC) on a Superdex 30 Increase 3.2/300 column (Cytiva) as described previously (Leitner et al., 2012).

Four fractions collected from the SEC step were analyzed in duplicate by liquid chromatography tandem mass spectrometry (LC-MS/MS) on a set-up consisting of an Easy nLC-1200 and an Orbitrap Fusion Lumos (both ThermoFisher Scientific). 5 to 10% of each SEC fraction were injected and separated on an Acclaim PepMap RSLC C_18_ column (250 mm × 75 µm, 2 µm particle size) at a flow rate of 300 nl/min. The mobile phases were A = water/acetonitrile/formic acid (98:2:0.15, v/v/v) and B = acetonitrile/water/formic acid (80:20:0.15, v/v/v), and the gradient ranged from 11 to 40 %B in 60 minutes.

MS data was acquired in the data-dependent acquisition / top speed mode with a cycle time of 3 s. A full scan in the orbitrap detected at 120 000 resolution was followed by data-dependent MS/MS scans at 30 000 resolution for precursors with a charge state between +3 and +7. Collision-induced dissociation was performed in the linear ion trap at a normalized collision energy of 35%. Dynamic exclusion was enabled for 30 s after one sequencing event.

Data analysis was performed using xQuest (Walzthoeni et al., 2012), version 2.5.1 (available from https://gitlab.ethz.ch/leitner_lab/xquest_xprophet). MS/MS spectra were searched against a database consisting of the sequences of the target protein constructs and three abundant contaminant proteins identified in a preliminary search. A decoy database for false discovery rate (FDR) control was generated by reversal and shuffling of the sequences using the xdecoy.pl script of xQuest. Search parameters included: Enzyme = trypsin, maximum number of missed cleavages = 2, peptide length = 4 to 40 amino acids, fixed modification = carbamidomethylation on Cys, variable modification = oxidation on Met, cross-linking sites = Lys and protein N terminus, cross-link mass shift = 138.06808 Da, difference between light and heavy linker = 12.07532 Da, MS mass error tolerance = ± 10 ppm, MS/MS mass error tolerance = ± 10 ppm. Post-search, the MS mass error window was adjusted to the actual offset ± 4 ppm, filtered according to a minimum xQuest score of 25, a TIC value of ≥0.1, a dSc value of ≤0.9, and manually evaluated. No decoy hits remained after this procedure, so that the FDR can be estimated as < ∼1% for intra- and < ∼5% for inter-protein cross-links.

## Cryo-EM Data collection

Data acquisition was performed on a Titan Krios (Thermo Fisher Scientific) operating at 300 kV equipped with a Gatan Imaging Filter (GIF) with a 20 eV energy slit using Gatan’s K3 direct electron detector in counting mode. Movies were collected using EPU software (Thermo Fisher Scientific) at a magnification of 130,000 and pixel size of 0.65 Å/pixel at a dose rate of approximately 8 e/pixel/s and a total dose of ca. 55 – 66 e/Å^2^ with defocus range of 0.5 to 2.5 µm.

## Cryo-EM image processing and model building

Image processing was carried out in cryoSPARC (Structura Biotechnology, Toronto, Canada) as outlined in the scheme (Supp. Figure 1). After drift correction, gain correction and averaging with Patch motion Correction, CTF fitting was performed with Patch CTF. Particles were selected on denoised micrographs using template picking.

2D class averages calculated by 2D classification of a subset of the sample selected by blob picking was used as template. After extraction (320 px box size binned to 96 px), 19844119 particles were subjected to 2D classification, retaining 3758671 particles, which were subsequently extracted without binning. Ab-initio model generation (10 classes) followed by heterogeneous refinement resulted in one class of 786052 particles that clearly contained DDB1, CUL4B, and additional density. Analysis of the class by heterogeneous refinement and subsequent 2D classification (70298 particles retained) revealed that all classes contained DDB1 and CUL4B, but only a few classes contained extra density corresponding to LIS1. Selection of 2D classes displaying additional density corresponding to LIS1 resulted after non-uniform refinement in a map with a resolution of 6.5 Å. Inspection of the map revealed conformational heterogeneity. After subjecting the particles to another round of 2D classification only 54950 particles exhibiting a clearly structured LIS1 density were retained, HR-HAIR (Kim et al., 2025) followed by non-uniform refinement resulted in a map with clear density for LIS1. Non-uniform refinement with a mask around DDB1 and LIS1 further improved the quality of the reconstruction to a resolution of 4.3 Å. The resolution of the LIS1 density is ∼8 Å, likely due to inherent flexibility, and therefore insufficient for building an atomic model. In order to determine the orientation of LIS1 with respect to DDB1 and analyze the interface between DDB1 and LIS1, we used Phenix ‘Dock in Map’ (Liebschner et al., 2019) to place the Alphafold3 (Jumper et al., 2021) model of LIS1 into the map. Figures were generated with PyMol (The PyMOL Molecular Graphics System, Version 3.0 Schrödinger, LLC.) and UCSF ChimeraX (Meng et al., 2023).

## Data and code availability

The cross-linking mass spectrometry-based proteomics data have been deposited to the ProteomeXchange Consortium via the PRIDE partner repository with the dataset identifier: PXD074695.

The AP-MS proteomics data have also been deposited to the PRIDE repository with the identifier: PXD074312

## Acknowledgement

We thank Anna Maria Stier, Claudia Schmidt, Piero Quattrocci, Martin Winkler and members of the Peter lab for helpful discussions, and Alicia Smith for critical editing. We thank Julia Kowal for her generous advice on cryo-EM data analysis. We acknowledge ScopeM and Miroslav Peterek for microscopy training and advise, and Ino Karemaker for help with the MS-analysis. Figure illustrations were prepared using Biorender (https://BioRender.com/r6ojq1v). We are grateful to D-BIOL for support by its maternity leave program. This work was financed by the Swiss National Science Foundation and ETH Zürich.

## Author contributions

W.I.W. and M.P. conceptualized the study. W.I.M performed the biochemical assays. W.I.M. and A.K. performed the AP-MS assay and F.U. analyzed the AP-MS data. W.I.M prepared the specimens for EM data collection, W.I.M. and B.M.Q imaged EM grids, and W.I.M., B.M.Q, and J.R. processed the EM data. A.L. performed the XL-MS analysis. M.P. and W.I.M. wrote the manuscript, with critical input from all authors.

---

**Supplementary Figure 1.** : Single-particle cryo-EM image processing workflow. Workflow and schematic representation of cryo-EM image processing, including a representative micrograph (scale bar: 30nm). The resolution map and FSC curve of the CUL4B-DDB1-LIS1 complex was locally refined with a focus on the DDB1-LIS1 interface. The resolution was determined using the 0.143 gold standard (Rosenthal & Henderson, 2003). Extended and closed conformations of CUL4B-DDB1-LIS1 complexes are marked with an asterisk.

**Supplementary Figure 2.** : Cryo-EM maps of the CUL4B-DDB1-LIS1 complex in the extended and closed state. **A.** Refined cryo-EM maps of the CUL4B-DDB1-LIS1 complex in the extended (left) and closed (right) conformation are shown in surface representation. The maps have been aligned to DDB1 (purple) and show the density arising from LIS1 (green) and CUL4B (grey). Additional density (white), potentially corresponding to the HLH motif, is observed but cannot be assigned with high confidence due to the low resolution (ca. 10 Å) of the maps. **B.** Crosslinking-mass-spectrometry analysis of 150 μg recombinant CRL4B^LIS1^ complexes incubated with 0.25 mM disuccinimidyl suberate (DSS) crosslinker. Cross-links among different subunits are indicated by green lines, and cross-links within the same subunit by purple lines. Note that CUL4B and LIS1 are fused, so they are considered as the same protein. The cross-links between CUL4B (CTD) and LIS1 (WD40) are labeled with yellow stars. The different subunits with domain boundaries are colored as follows: DDB1 in purple, LIS1 in green, CUL4B in gray, and Rbx1 in blue.

**Supplementary Figure 3.** : REKE loop mutations decrease LIS1 binding to the CRL4 complex. **A.** Size exclusion elution profiles (280 nm) of the last purification step of recombinantly expressed CUL4B-RBX1-DDB1 (WT) and CUL4B-RBX1-DDB1 mutant (EAAA) complexes, and biotinylated LIS1 and DCAF8, respectively. **B.** Fluorescent scans and Coomassie-stained SDS–PAGE of *in vitro* FLAG pull-down assays using FLAG-tagged LIS1 (F-LIS1), Strep-tagged CUL4B (S-CUL4B) and Strep-tagged wild-type (S-DDB1) and either EAAA or AAAA mutant DDB1, co-expressed in baculoviral Hi5 cells. One of three biological replicates is shown. The band intensities of the fluorescent bands of wild-type (WT) and the indicated DDB1 mutants (EAAA and AAAA) (right) was quantified and normalized to the input. **C.** Western blot analysis of *in vitro* Strep pull-downs using purified components of 0.25 μM Strep-tagged CUL4B-RBX1-DDB1 (WT) and CUL4B-RBX1-DDB1 (EAAA) mutant complexes and increasing concentrations of untagged LIS1 (at 0, 0.2, 0.8, 3, and 5 μM) (left). Quantification of the chemiluminescence HRP signal corresponding to LIS1 (right). Note that binding of LIS1 to the DDB1-EAAA mutant complex (blue bars) is reduced compared to the wild-type control (violet bars). **D.** Coomassie-stained SDS–PAGE of *in vitro* Strep (S) pull-down of His (H)-tagged DDB1 (WT) or mutant (AAAA) and Strep-DCAF8 co-expressed in Hi5 cells. **E.** Conservation of the DDB1 REKE loop across different species was assessed using Jalview (Waterhouse et al., 2009) with ClustalW (Larkin et al., 2007) for multiple-sequence alignment. The color-code highlights conservation of critical residues. **F.** Close-up view of the cryo-EM structure of DDB1(ΔBPB)-DDA1-DCAF5 at 2.63 Å (PDB: 8TL6) showing the helix of DDA1 approaching the REKE loop of DDB1. The distance between DDB1 (E201) and DDA1 (R57) is 3.5 Å. **G.** Close-up view of the crystal structure of DDB1-DDA1-DCAF15-RBM39 at 2.9 Å (PDB: 6PAI) showing the DDA1 helix approaching the REKE loop of DDB1. Distances are specified as follows: DDB1 (E201) and DDA1 (R57) 4.5 Å and DDB1 (E199) and DDA1 (K65) 5.1 Å.

**Supplementary Figure 4.** : Phosphorylation of the CUL4B N-terminal domain does not affect DDB1 incorporation into the CRL4B complex. **A.** Size exclusion elution profile (280 nm) of the final purification of recombinantly expressed CUL4B-RBX1. The dashed box highlights the complex. **B.** Coomassie-stained SDS–PAGE of recombinantly expressed CUL4B-RBX1 complex, before and after treatment with λ-Protein Phosphatase for 2 hours. The position of phosphorylated (P-CUL4B) and unphosphorylated CUL4B is indicated. **D.** Schematic diagram of FP assays measuring the binding affinity of phosphorylated or dephosphorylated (λ-PP treated) CUL4B-RBX1 complexes to Alexa488 (orange)-labeled DDB1 (purple). FP binding curves were measured using increasing concentrations of phosphorylated P-CUL4B-RBX1 (gray curve) or dephosphorylated CUL4B-RBX1 (black curve) in the presence of a fixed concentration of DDB1. Quantified Kds (μM) are listed in the table. Note that CUL4B phosphorylation does not regulate DDB1 binding.

**Supplementary Figure 5.** : Identification of CUL4B-specific DCAFs that require its N-terminal extension and validation of BRWD1. **A.** Abundance of the bait recovery (CUL4B) expressed as spectra counts across the following conditions: control without tag, CUL4B and CUL4B ΔN. **B.** Boxplot showing the log2 intensity of proteins across all replicates for co-immunoprecipitation control, CUL4B and CUL4B ΔN **C.** Volcano plot showing the log2 fold change enrichment of proteins co-immunopurified with CUL4B and CUL4B ΔN (respectively left and right). CUL4B interactors annotated in the volcano plot were enriched with a log2FC> 1 and pvalue < 0.05 against the control purification (purification without tag). The significance was calculated as two-sided unpaired Students’s t-test from three biological replicates. **D.** Close-up view of AlphaFold 3 structural prediction of DDB1 (Δ-BPB) bound to BRWD1 (Δ-1451-2182). BRWD1 uses an extended helix that contacts the DDB1 REKE loop in the BPA domain. The interaction surface involves similar residues in DDB1 (BPA) as those used for binding LIS1. Distances are specified as follows: DDB1 (E201) and BRWD1 (R775) 5.9 Å, DDB1 (R158) and BRWD1 (E398) 6.7 Å, DDB1 (R111) and BRWD1 (E760) 3.8 Å. **E.** SDS-PAGE gel of *in vitro* reconstituted Strep (S)-tagged CUL4B-DDB1 complexes with either wild-type (WT) DDB1 or the REKE mutant (EAAA). 5.3 µM of these purified complexes (Strep-pull down) were incubated with lysates of Hi5 cells over-expressing GST-tagged BRWD1. Bound BRWD1 was visualized by immunoblotting with an anti-BRWD1 antibody and quantified using a fluorescent secondary antibody (bottom). The presence of DDB1 and CUL4B was visualized by the stain-free BioRad system.

