## Supplementary figures for "Molecular determinants underlying substrate receptor specificity of human CRL4B E3 ubiquitin ligase"

**Supplementary Figure 1**

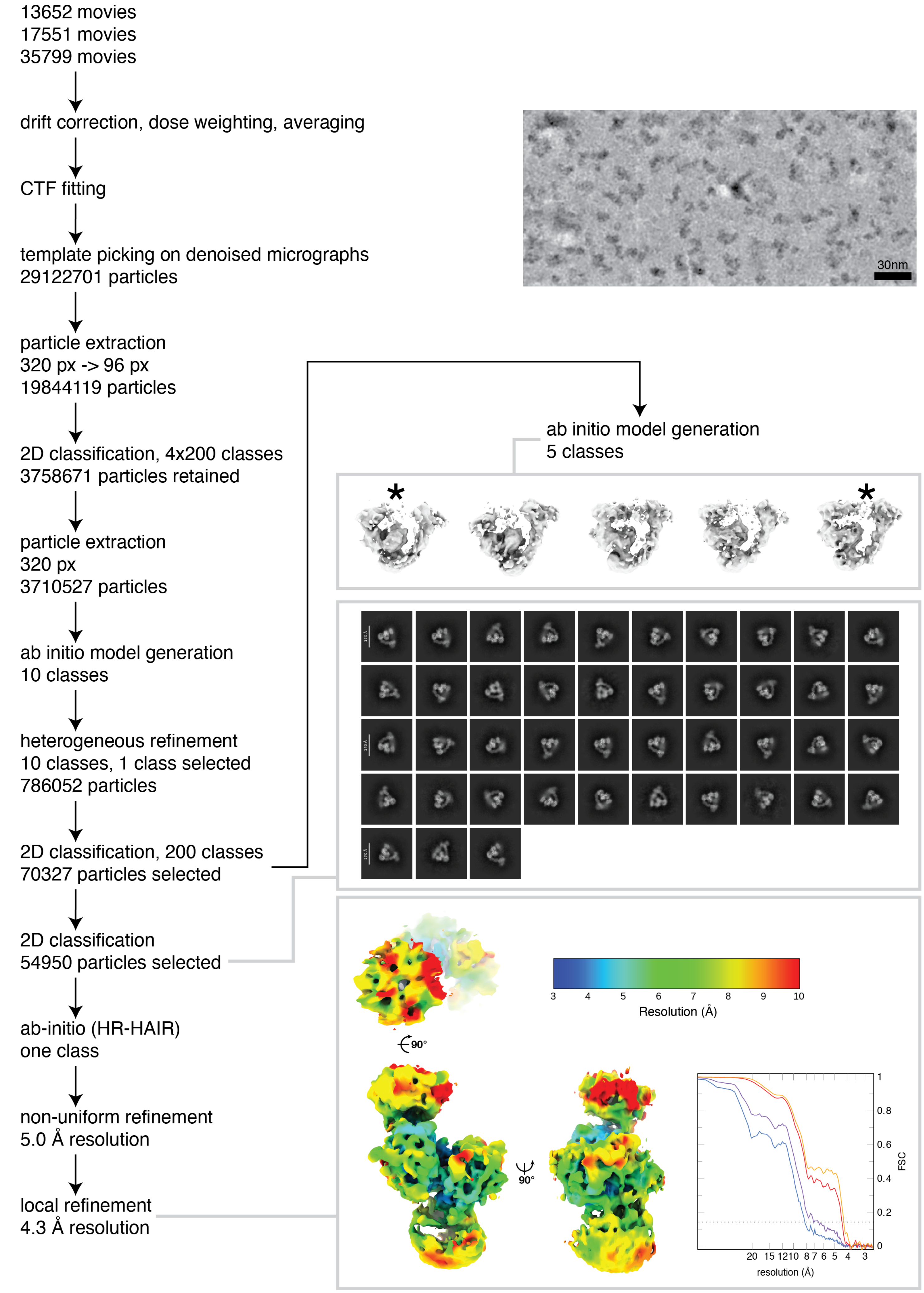

Supplementary Figure 2

**A.**

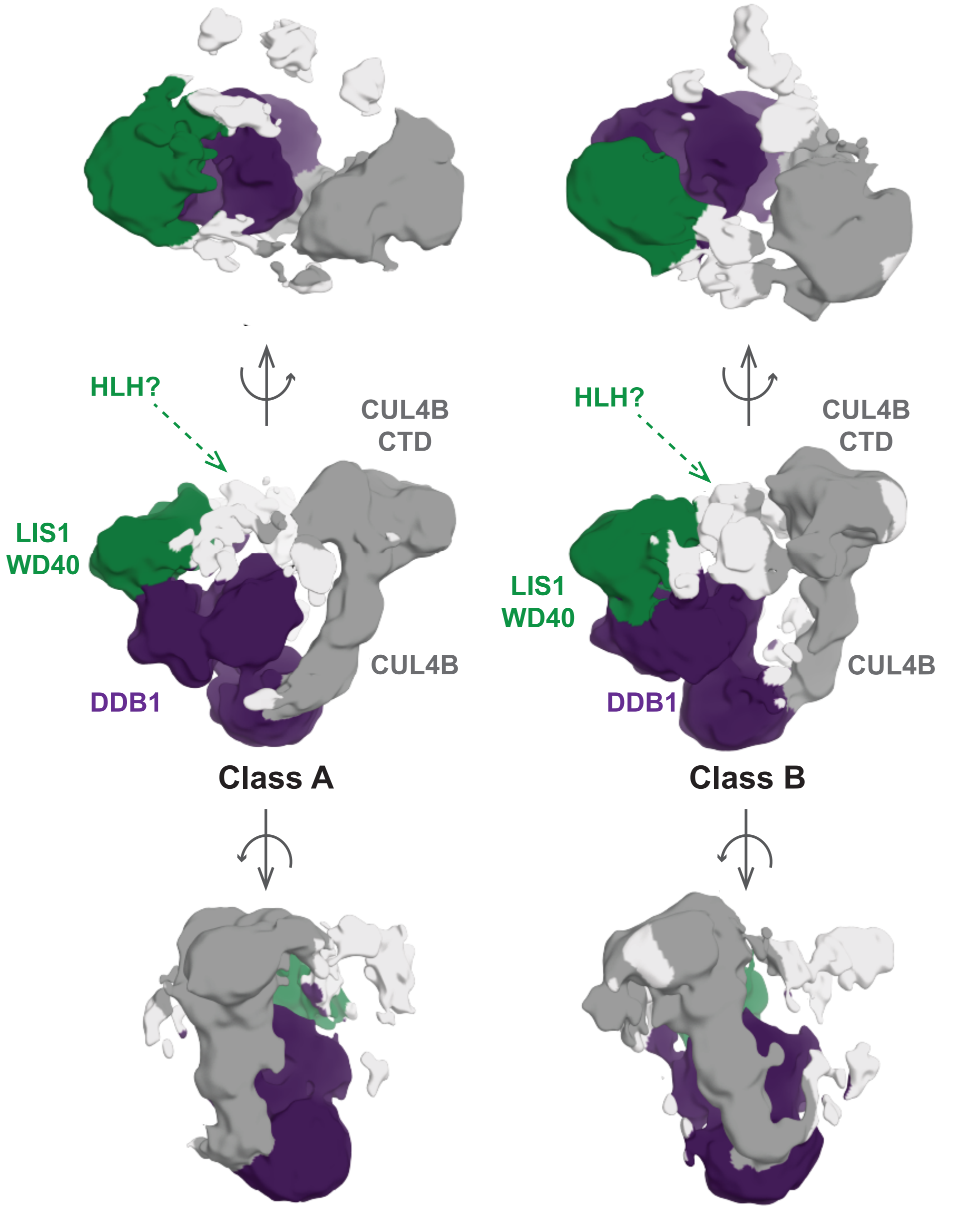


**B.**

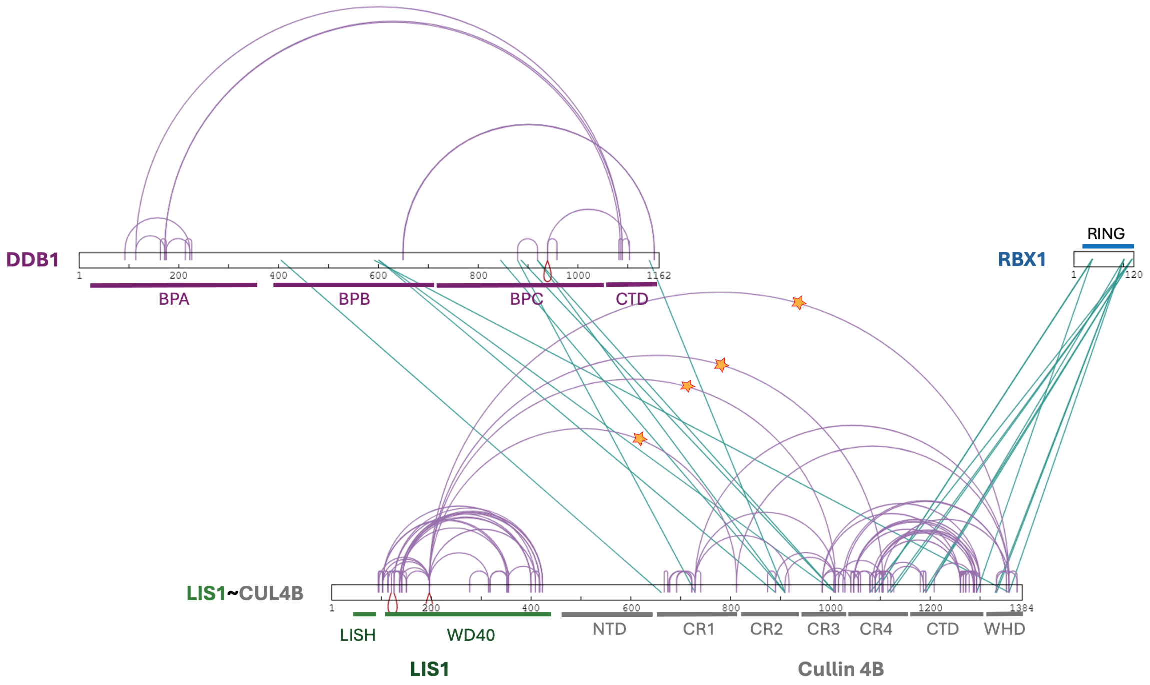

#### Supplementary Figure 3

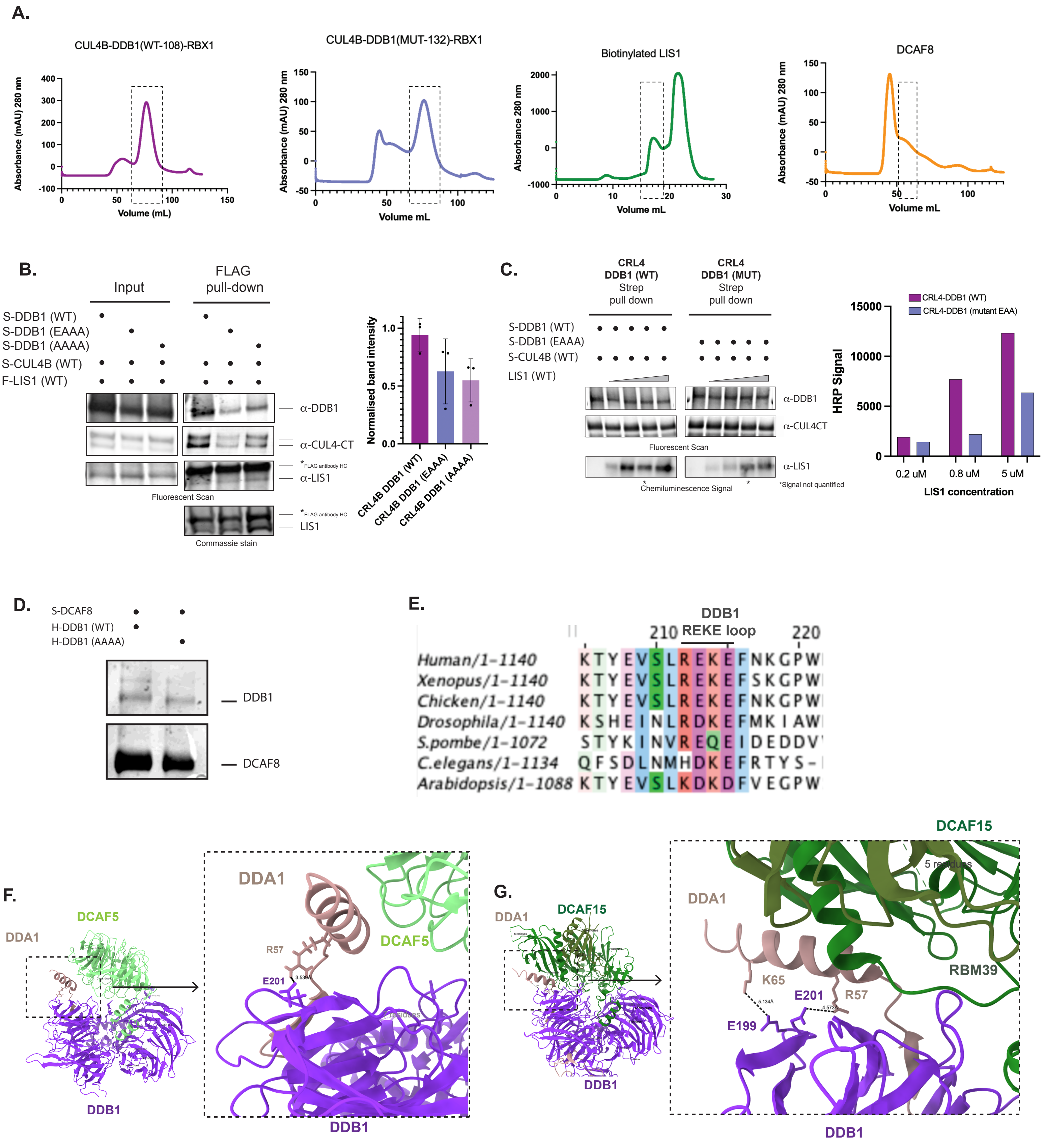

### Supplementary Figure 4

A.

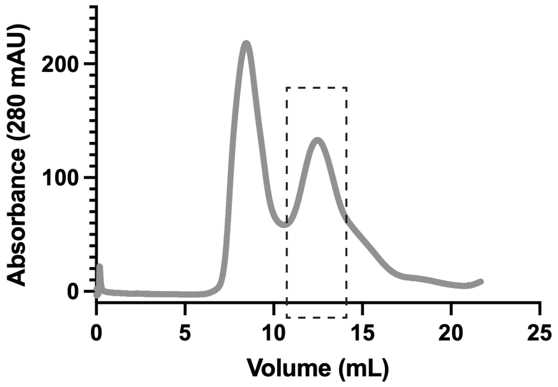


B.

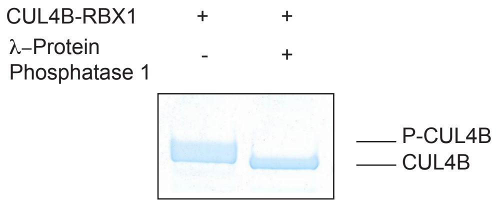


C.

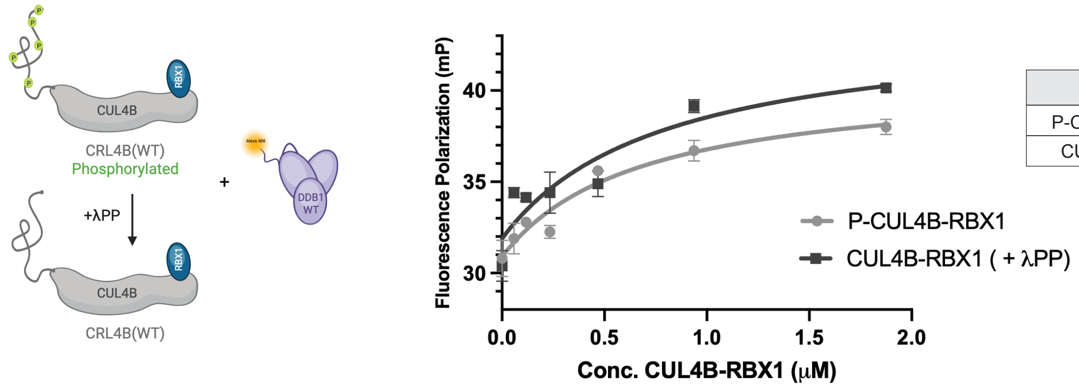


| Protein | $K_d$ (μM) |
|---|---|
| P-CUL4B-RBX1 | 0.67 |
| CUL4B-RBX1 | 0.78 |

Supplementary Figure 5

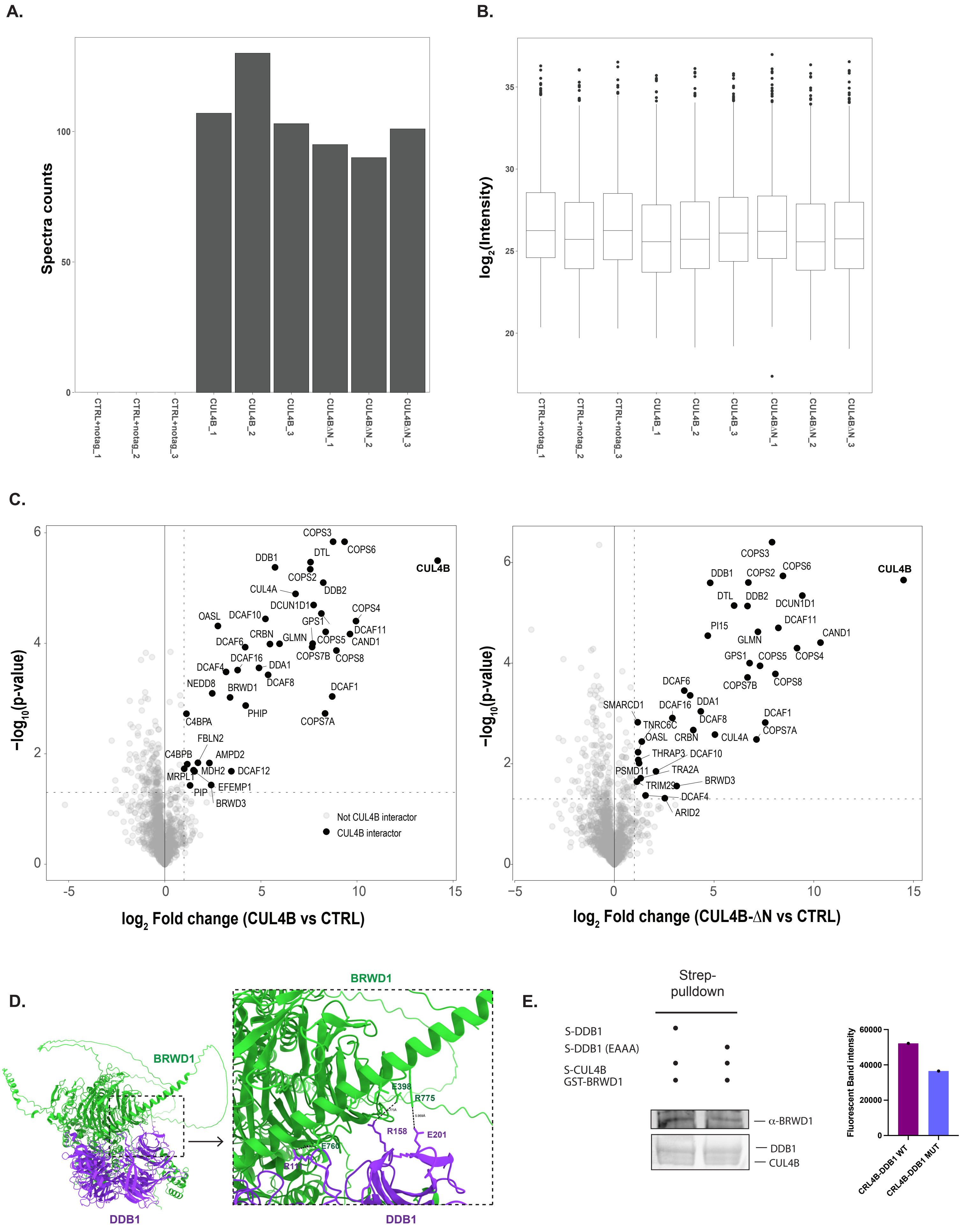
